# *C. elegans* Runx/CBFβ suppresses POP-1(TCF) to convert asymmetric to proliferative division of stem cell-like seam cells

**DOI:** 10.1101/625335

**Authors:** Suzanne E. M. van der Horst, Janine Cravo, Alison Woollard, Juliane Teapal, Sander van den Heuvel

## Abstract

A correct balance between proliferative and asymmetric cell divisions underlies normal development, stem cell maintenance and tissue homeostasis. What determines whether cells undergo symmetric or asymmetric cell division is poorly understood. To gain insight in the mechanisms involved, we studied the stem cell-like seam cells in the *Caenorhabditis elegans* epidermis. Seam cells go through a reproducible pattern of asymmetric divisions, instructed by non-canonical Wnt/β-catenin asymmetry signaling, and symmetric divisions that increase the seam cell number. Using time-lapse fluorescence microscopy, we show that symmetric cell divisions maintain the asymmetric localization of Wnt/β-catenin pathway components. Observations based on lineage-specific knockout and GFP-tagging of endogenous *pop-1* support the model that POP-1^TCF^ induces differentiation at a high nuclear level, while low nuclear POP-1 promotes seam cell self-renewal. Before symmetric division, the transcriptional regulator *rnt-1*^Runx^ and cofactor *bro-1*^CBFβ^ temporarily bypass Wnt/β-catenin asymmetry by downregulating *pop-1* expression. Thereby, RNT-1/BRO-1 appears to render POP-1 below the level required for its repressor function, which converts differentiation into self-renewal. Thus, opposition between the *C. elegans* Runx/CBFβ and TCF stem-cell regulators controls the switch between asymmetric and symmetric seam cell division.

## INTRODUCTION

Tissue-specific stem cells combine long-term maintenance with the generation of differentiating daughter cells. This can be achieved by asymmetric cell divisions that simultaneously generate a self-renewing stem cell and a daughter cell that initiates a differentiation program (Reviewed in Knoblich 2010). Expanding stem cell numbers, however, requires symmetric divisions that generate two self-renewing stem cells. Thus, the proper balance between symmetric and asymmetric divisions is key to the development and maintenance of tissues, and to preventing tumor formation or premature differentiation. How this balance is controlled is currently poorly understood.

The *Caenorhabditis elegans* epidermis provides an attractive model to study stem cell divisions in a developing tissue. The stem cell-like seam cells form part of the epidermis and undergo a reproducible pattern of symmetric and asymmetric divisions at stereotypical times of development (Sulston & Horvitz 1977). Asymmetric divisions of seam cells create a new seam daughter cell, as well as a cell that proceeds either to form neurons or to differentiate and fuse with the general epidermis (known as hypodermis in *C. elegans*). In addition, the number of seam cells increases in the second larval stage (L2), through symmetric divisions that generate two seam daughter cells.

Non-canonical Wnt signaling, mediated by the Wnt/β-catenin asymmetry pathway, is critical for many asymmetric cell divisions in *C. elegans*, including seam cell divisions (Lin et al. 1998; Kidd et al. 2005; Mizumoto & Sawa 2007a; Baldwin & Phillips 2014). This pathway controls the choice between two alternative cell fates, instructed by an unequal subcellular localization of Wnt/β-catenin pathway components. Ultimately, the different cell fates are determined by asymmetric activity of the TCF/LEF-related transcription factor POP-1 (posterior pharynx defective) (Lin et al. 1998). POP-1 is thought to function as a transcriptional repressor in a complex with UNC-37 (Groucho) that induces differentiation (Calvo et al. 2002). POP-1 can also function as a transcriptional activator with co-factor SYS-1 (β-catenin) instructing self-renewal (Kidd et al. 2005; Shetty et al. 2005; Huang et al. 2007). Wnt signaling and asymmetric localization of upstream pathway components restrict the repressor function to anterior cells, through export of POP-1 from the nucleus of posterior seam daughter cells (Takeshita & Sawa 2005), and by degrading the co-activator SYS-1 in the differentiating anterior daughters (Vora & Phillips 2015). Altered levels or localization defects of several Wnt/β-catenin asymmetry pathway components result in symmetric seam cell divisions, indicating the importance of this pathway for division asymmetry (Banerjee et al. 2010; Gleason & Eisenmann 2010; Ren & Zhang 2010; Hughes et al. 2013). Whether and how symmetric seam cell divisions circumvent the Wnt/β-catenin asymmetry pathway is currently not understood.

A conserved Runx transcriptional repressor complex also contributes to the control of seam cell division and differentiation. Runx transcription factors play broad functions in development and stem cell maintenance, and are probably best known for their critical roles in hematopoiesis and oncogenic functions in leukemia (Reviewed in Deltcheva & Nimmo 2017). They act in association with a heterodimeric partner, CBFβ, and contribute to repression as well as activation of transcription. The *C. elegans* genome encodes a single Runx homolog, RNT-1, and single CBFβ-related cofactor, BRO-1 (Nimmo et al. 2005; Kagoshima et al. 2007a; Xia et al. 2007). Genetic and biochemical experiments support that RNT-1 and BRO-1 form a transcriptional repressor complex together with UNC-37^Groucho^. Mutations in *rnt-1, bro-1* and *unc-37* reduce the seam cell number as a consequence of defects in the L2 division pattern (Nimmo et al. 2005; Kagoshima et al. 2007a; Xia et al. 2007). By contrast, induced expression of RNT-1 and BRO-1 increases the seam cell number. These observations highlight a regulatory role for the RNT-1/BRO-1 complex in seam cell proliferation and differentiation. It remains unclear, however, how this is integrated with Wnt/β-catenin asymmetry signaling to establish the reproducible pattern of symmetric and asymmetric seam cell divisions, and previous studies concluded that these regulators act in parallel (Kagoshima et al. 2005; Gleason & Eisenmann 2010; Hughes et al. 2013).

In this study, we use time-lapse fluorescence microscopy of developing larvae to identify the mechanisms that determine asymmetric versus proliferative seam cell division. We show that anterior daughter cells adopt a seam cell fate during symmetric cell divisions despite asymmetric distribution of Wnt/β-catenin asymmetry pathway components. This indicates that symmetric divisions bypass Wnt/β-catenin asymmetry to prevent anterior cell differentiation. Multiple observations support that the RNT-1/BRO-1 complex provides this bypass-mechanism by temporarily repressing *pop-1*. First, GFP-tagged endogenous POP-1 is expressed at a very low level during symmetric seam cell divisions, dependent on *rnt-1 bro-1* function. Further, double *rnt-1 bro-1* mutants show ectopic differentiation of anterior seam cells, which is fully suppressed by *pop-1* RNAi. Moreover, induced expression of RNT-1/BRO-1 represses GFP::POP-1 expression and turns asymmetric seam cell divisions into symmetric divisions. Finally, endogenous RNT-1 is expressed at a high level before symmetric seam cell divisions, but disappears and remains absent prior to the subsequent asymmetric division, which correlates with upregulation of POP-1. These data support the model that RNT-1/BRO-1 provides temporal control over POP-1^TCF/LEF^, which renders POP-1 below a critical level required for its repressor function, and thereby changes differentiation into self-renewal. Together, our data reveal how interactions between two conserved stem cell regulators can balance symmetric and asymmetric divisions in a developing tissue.

## RESULTS

### Wnt components localize asymmetrically in symmetric seam cell divisions

We studied the stem cell-like precursors of the *C. elegans* epidermis to reveal the mechanisms that determine whether cells undergo symmetric or asymmetric cell divisions. The seam cells reside in two lateral epithelia along the anterior-posterior body axis (Fig. 1). During the first larval stage, each V seam cell undergoes one anterior-posterior oriented asymmetric division (Sulston and Horvitz, 1977). These divisions generate a self-renewing posterior daughter cell and an anterior daughter cell that either differentiates and fuses with the epidermis (V1-V4, V6) or forms neuronal daughter cells (V5). Upon entry of the second larval stage (L2), V1-V4 and V6 go through a symmetric division to generate two self-renewing seam daughter cells. This symmetric division is followed by an asymmetric division of the V cells to produce epidermal (V1-V4, V6) and neuronal (V5) cells.

**Figure 1.**
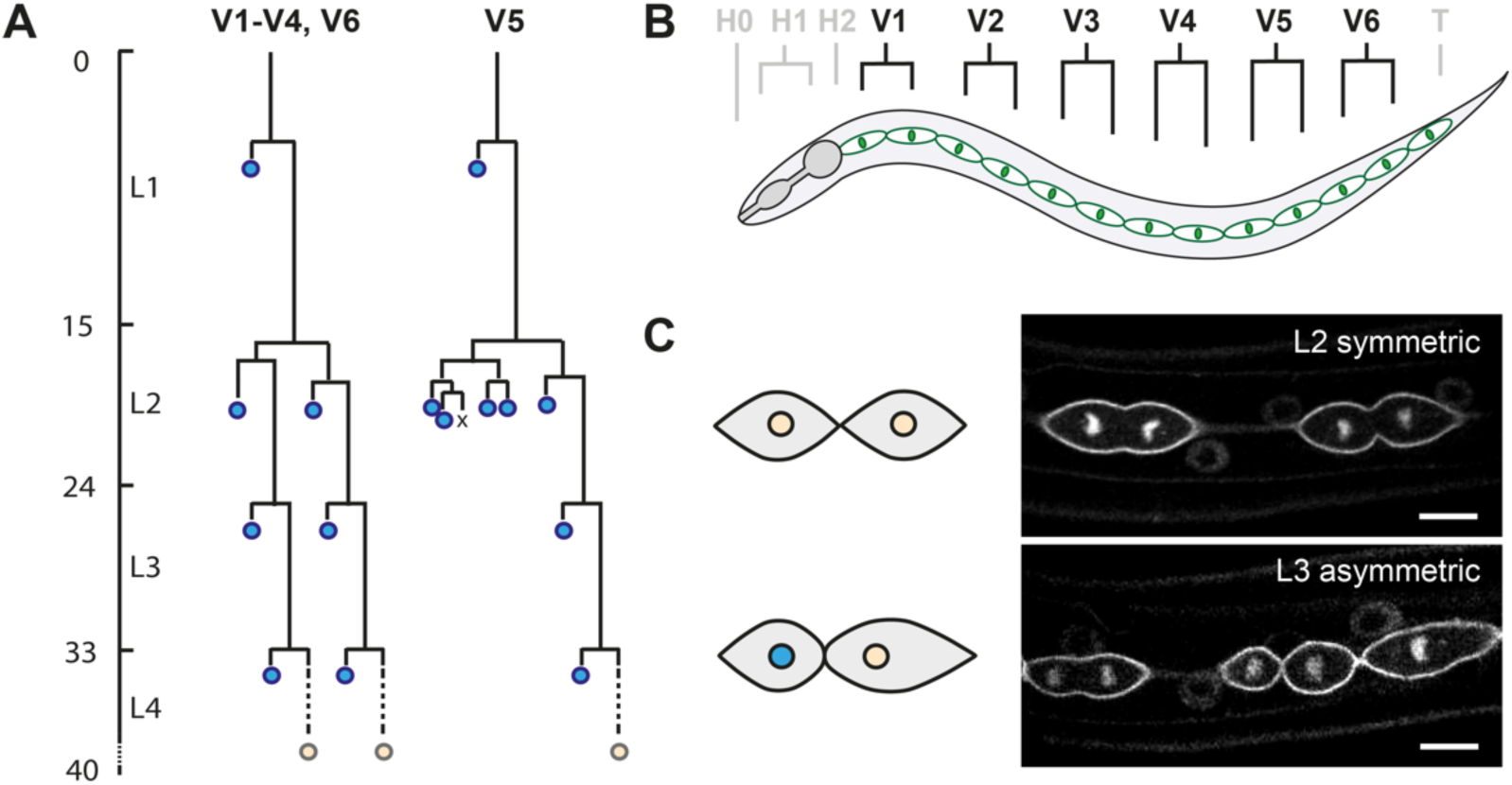
Seam cell lineage as a model to study the regulation of proliferative versus asymmetric cell division. (A) Postembryonic division patterns of the ventrolateral precursor (V) cells of the *C. elegans* epidermis (hypodermis). The seam cells undergo cell division (horizontal lines) in a stereotypic manner during each of the four larval stages (L1-L4), as indicated by the time course of development (left axis). Asymmetric divisions of V1-V4 and V6 generate one anterior epidermal daughter cell (blue), and one self-renewing posterior seam daughter cell. At the end of larval development, all remaining seam cells (orange) exit the cell cycle and fuse together to form two lateral syncytia. (B) Schematic lateral view of one of two seam epithelia. The anterior region includes the epidermal precursor cells of the head (H0-H2), the middle ventrolateral region contains the V cells (V1-V6) and the posterior region hosts the tail blast cell (T). (C) Representative spinning disk confocal fluorescence microscopy images of seam cells expressing transgenic reporter genes, to visualize the membrane (GFP::PH) and DNA (GFP::H2B). The L2 symmetric divisions generate two self-renewing daughter cells equal in cell size and fate (orange nucleus; top). The asymmetric division generates a smaller anterior daughter cell that will fuse with the epidermis (blue nucleus), and a larger posterior self-renewing seam cell (orange nucleus; bottom). Scale bars represent 10 µm.

We examined the distribution of Wnt/β-catenin asymmetry pathway components to gain insight in the regulation of symmetric versus asymmetric seam cell division. Earlier observations indicated that the asymmetric localization of POP-1 and APR-1 is maintained during the symmetric divisions of seam cells in L2 (Wildwater et al. 2011; Baldwin & Phillips 2014). To follow this process more closely, we made use of spinning disk time-lapse fluorescence microscopy and the P*sys-1::pop-1::gfp* reporter. This transgene was previously used to demonstrate unequal nuclear POP-1 levels during the asymmetric divisions of V5 and T cells (Kagoshima et al. 2005; Mizumoto & Sawa 2007b). We observed a similar pattern of POP-1 localization during the asymmetric divisions of seam cells in the V1-4, V6 lineages (Fig. 2A, bottom). Soon after the nuclei reformed in telophase, POP-1::GFP levels decreased in the posterior nucleus, in contrast to the anterior nucleus. Quantification of the fluorescence intensity indicated an approximately 2-fold nuclear enrichment of POP-1 in the anterior compared to posterior daughter cells at the time of cytokinesis (Fig. 2B).

**Figure 2.**
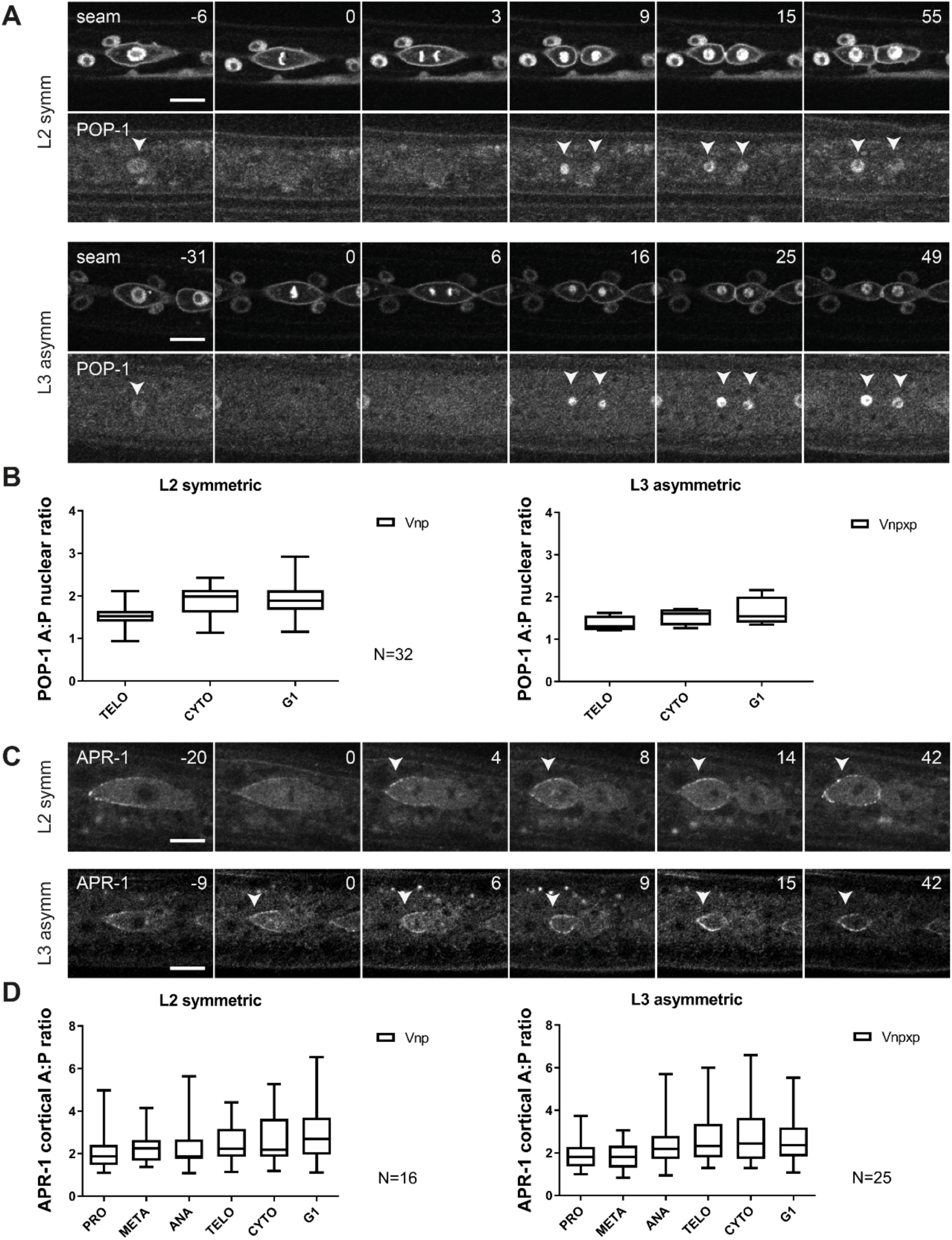
Localization dynamics of POP-1 and APR-1 during seam cell divisions. (A) Representative images from spinning disk time-lapse microscopy, showing seam markers mCherry::PH and mCherry::H2B, and POP-1::GFP during L2 symmetric (upper panels) and L3 asymmetric divisions (bottom panels). Arrowheads point to seam cell nuclei. Anterior is to the left. (B) Quantification of the POP-1 nuclear A:P ratio in L2 symmetric (left) and L3 asymmetric divisions (right). (C) Images from spinning disk time-lapse microscopy of APR-1::VENUS during L2 symmetric (upper panel) and L3 asymmetric divisions (bottom panel). Arrowheads point to the anterior cortex of a dividing seam cell. Anterior is to the left. (D) Quantification of the APR-1 cortical A:P ratio in L2 symmetric (left) and L3 asymmetric divisions (right). The box and whiskers plots indicate mean (line within box) as well as the highest and lowest observed values (lines outside box). Images were processed using ImageJ software, the scale bar (10 μm) is the same for all images.

The lower nuclear level of POP-1 was previously shown to correspond to activation of the Wnt pathway and acquisition of the seam cell fate (Gleason & Eisenmann 2010; Gorrepati et al. 2013). Notably, however, the L2 symmetric divisions that generate two seam daughter cells also showed asymmetric POP-1 distribution. During telophase and cytokinesis of symmetric seam cell divisions, the POP-1::GFP levels decreased specifically in the posterior nucleus (Fig. 2A, top). These live observations confirm our previous conclusion based on immunohistochemical detection of POP-1::GFP (Wildwater et al. 2011), and indicate that the mechanisms for asymmetric distribution of POP-1 remain active during the symmetric divisions that create two seam daughter cells.

To examine this aspect further, we followed the localization of APR-1, making use of a P*apr-1::apr-1::venus* reporter (Mizumoto & Sawa 2007a). As described before, APR-1 enriches at the anterior half of the cell cortex of asymmetrically dividing seam cells, and upon completion of cytokinesis is predominantly detected at the cortex of anterior daughter cells (Fig. 2C). Similar to the asymmetric L3 divisions, we observed anterior enrichment of APR-1 during the L2 symmetric divisions (Fig. 2C). Quantifications of APR-1::VENUS levels confirmed that the ratio’s between anterior versus posterior cortical levels of APR-1 were similar between symmetric and asymmetric cell divisions (Fig. 2D). Together, these live observations confirm that anterior-posterior polarization of Wnt/β-catenin asymmetry components also takes place during symmetric divisions. Despite this asymmetric APR-1 and POP-1 distribution, the anterior daughter cells do not differentiate but adopt a seam cell fate that is normally restricted to the posterior daughter cell. This appears to imply that the Wnt/β-catenin asymmetry pathway is temporarily overruled during symmetric seam cell divisions.

### Continued proliferation does not overrule the Wnt/β-catenin asymmetry pathway

Cell cycle progression and CDK-cyclin activity are generally considered to oppose cell differentiation (Ruijtenberg & van den Heuvel 2016). In contrast to asymmetric divisions, the symmetric seam cell divisions are rapidly followed by a second round of cell division (Sulston & Horvitz 1977). The localization of a cyclin-dependent kinase (CDK) sensor supports that both daughter cells of symmetric seam cell divisions immediately progress into the next cell cycle and fully activate CDKs (Fig. S1A) (Van Rijnberk et al. 2017). The anterior daughter cells initiate the next mitosis approximately 2 hours after the completion of symmetric cell divisions in L2. This falls within the time lag observed for the onset of differentiation in other larval stages, as defined by fusion of anterior daughter cells with the hypodermis (2-2,5 hours after asymmetric division). As differentiation normally coincides with low CDK activity, we wondered whether cell cycle progression and high CDK activity overrules POP-1/UNC-37 induced differentiation in anterior seam daughter cells.

To test this possibility, we examined whether inducing or arresting cell cycle progression changes the normal seam daughter cell pattern. Heat shock-induced expression of CDK-1, CYB-1 and CYB-3 just before asymmetric divisions in L2 or L3 occasionally induced extra cell division. The daughter cells of these divisions retained the anterior or posterior fate, continued with an extra asymmetric division, or fused with each other (Fig. S2). These induced divisions appeared abnormal, however, with cells maintaining condensed DNA, rapidly re-entering mitosis and possibly skipping S phase. To examine a more physiological situation, we arrested the cell cycle after symmetric cell division, using heat shock-induced expression of the CDK inhibitor *cki-1* CIP/KIP. Time-lapse fluorescence microscopy showed that this resulted in substantial CKI-1::GFP levels in seam cells (Fig. 3). Expression of CKI-1::GFP after the symmetric divisions in L2 suppressed the second round of seam cell divisions by more than 5 hours (Fig. 3A, bottom). The large majority of the arrested anterior seam daughter cells did not show signs of differentiation (88% of the Vn.ppa cells retained the seam fate, n =37). As a control for cell-cycle independent effects, we induced *cki-1::gfp* expression in seam daughter cells after the L1 asymmetric divisions. At this time, anterior seam daughter cells continued differentiation as normal, with 100% of Vn.a cells fusing with the epidermis (Fig. 3A, Top). Based on these observations, it appears unlikely that the rapid continuation of the next cell division cycle is the mechanism that overrules the Wnt/β-catenin asymmetry pathway in anterior seam cells.

**Figure 3.**
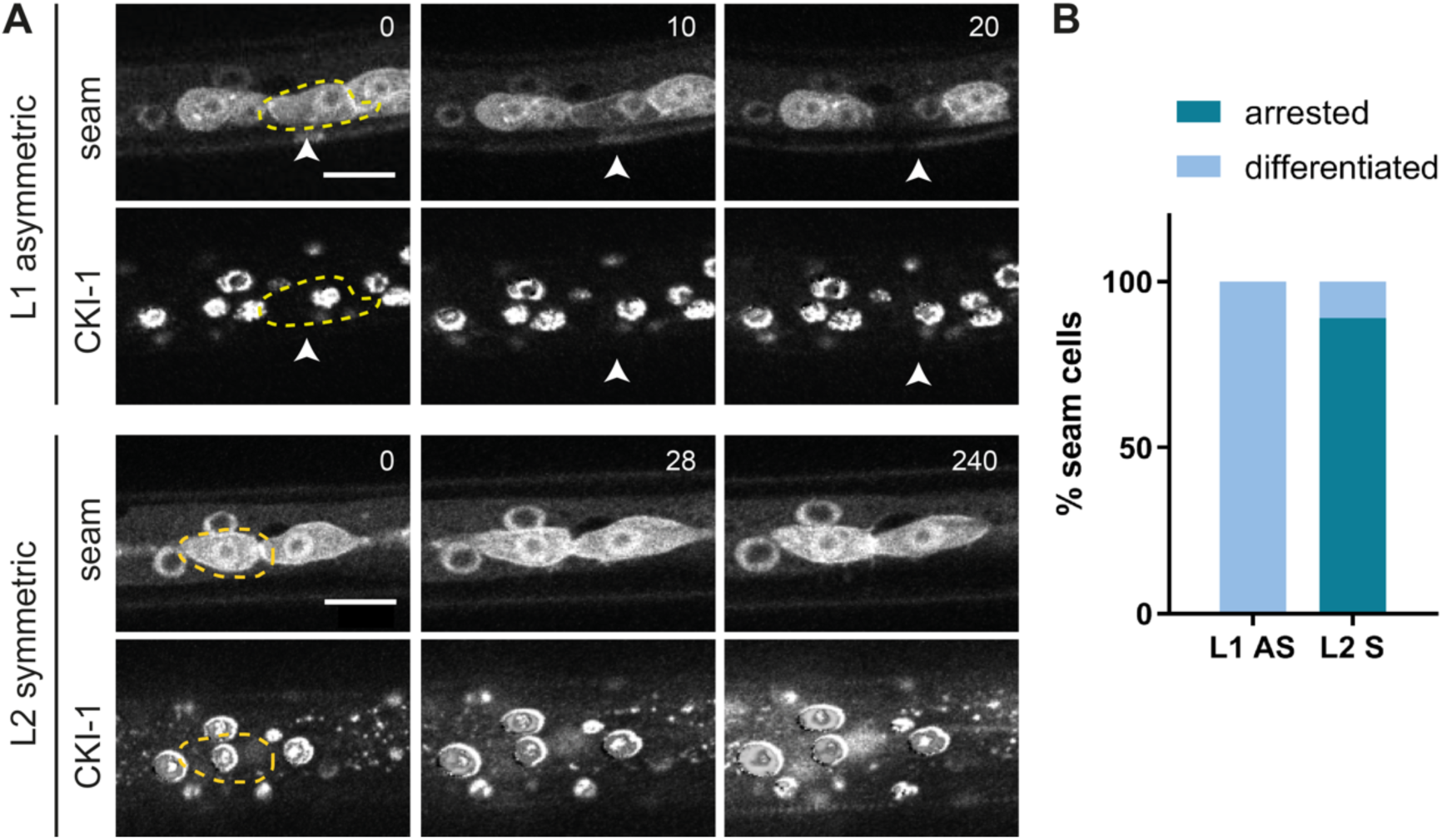
CKI-1 induction in L1 and L2 seam cells. (A) Time-lapse recording of P*hsp::cki-1::gfp* animals, heat-shocked around the time of L1 division (upper panels) or between symmetric and asymmetric divisions in L2 (bottom panels). Time series (minutes) started 1 hour after heat shock induction. Images show the seam markers mCherry::PH and mCherry::H2B (upper panels) and CKI-1::GFP (lower panels). Anterior daughter cells are outlined (yellow), the arrow heads indicate a differentiating Vn.a daughter cell. Scale bars represent 10 µm. (B) Quantification of the number of anterior daughter cells that differentiate after CKI-1 induction (grey) or that retain seam fate (black) in L1 and L2 animals.

### The RNT-1/BRO-1 transcriptional repressor complex promotes seam cell fate

Another candidate mechanism to overrule Wnt/β-catenin asymmetry is the RNT-1/BRO-1 transcriptional repressor complex. Studies of *rnt-1* and *bro-1* loss-of-function mutants revealed L2-specific seam cell division defects in hermaphrodites, as well as V6 and T division defects during the development of the male tail (Kagoshima et al. 2005, 2007b; Nimmo et al. 2005; Xia et al. 2007). Using spinning disk time-lapse microscopy, we followed L2 seam cell divisions in the candidate double null mutant *rnt-1(tm388) bro-1(tm1183)*. Control animals showed the reproducible pattern of one round of symmetric divisions followed by asymmetric cell divisions at the reported stereotypical times. The timing of L2 seam cell division was not altered in *rnt-1 bro-1* double mutant animals, but variable defects in the division pattern were observed. 74% of *rnt-1 bro-1* Vn.p seam cells skipped at least one cell division in L2 (43% skipped the posterior asymmetric division, and 31% skipped the symmetric division). In this latter group, the anterior Vn.pa daughter cell inappropriately underwent differentiation and fused with hyp7 (31% of the lineages; Fig. 4A). The posterior daughter remained a seam cell when the asymmetric division was omitted; hence this defect does not alter the seam cell number at later stages. The missed symmetric divisions reduced the seam cell number to approximately 13 per lateral side, compared to 16 in wild-type animals (Fig. 4B,C). The skipped divisions and immediate differentiation of anterior seam daughter cells in L2 confirm previous observations (Kagoshima et al. 2007a; Xia et al. 2007), and indicate that RNT-1 and BRO-1 normally act to both promote proliferation and prevent differentiation of seam cells.

**Figure 4.**
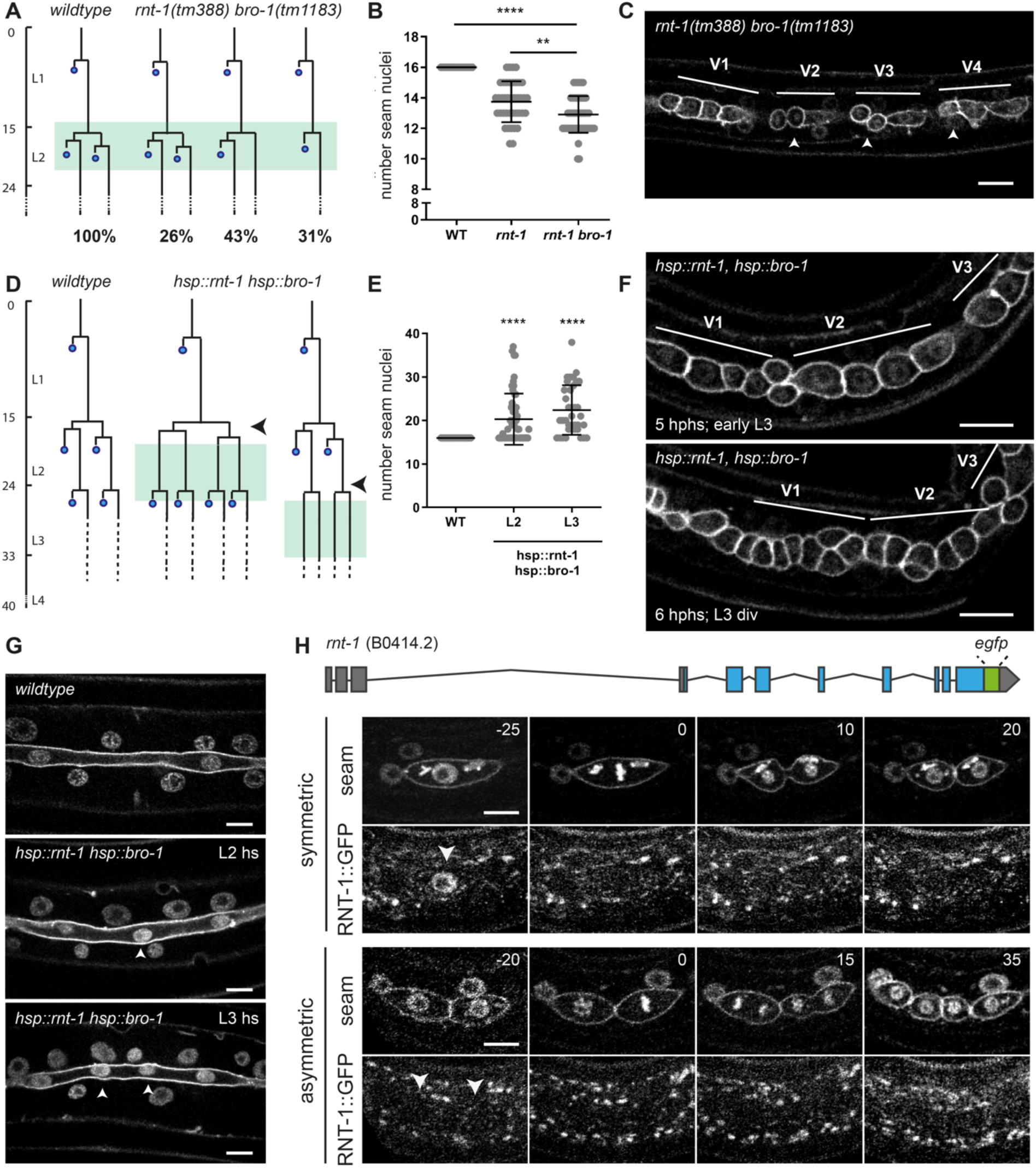
The RNT-1/BRO-1 transcriptional repressor complex promotes the seam cell fate. (A) Lineage analysis of L2 divisions of control animals and the *rnt-1(tm388) bro-1(tm1183)* double mutant (green box marks the time window of analysis). (B) Quantification of the number of seam cell nuclei at the end of L4 development for wild-type control animals, the single *rnt-1(tm388)* mutant and *rnt-1(tm388) bro-1(tm1183)* double mutant larvae. (C) Representative spinning disk confocal image of the seam cell epithelium of *rnt-1(tm388) bro-1(tm1183)* animals in the mid L2 stage. V1 division was normal (representing the 26% lineage in A), V2, V3 and V4 anterior daughter cells undergo the L2 asymmetric division (arrow heads), whereas the posterior cells do not (representing the 43% lineage in A) (D) Lineage analysis of control animals and heat-shock induced *rnt-1,bro-1* animals. Heat-shock was given between the symmetric and asymmetric division in L2 (middle; arrowhead) or prior to the L3 asymmetric division (right; arrowhead). Late L2 cells and L3 divisions were followed (green boxes). (E) Quantification of the number of seam nuclei at the end of L4 development for control animals and heat-shock-induced L2 and L3 animals. (F) Time-lapse spinning disk microscopy images of early L3 animals that underwent heat-shock induction of *rnt-1 bro-1* at t=17.30-18.30 hr, between the symmetric and asymmetric L2 divisions. Images show epithelium 5 hours after the end of heat-shock (top; early L3 t=23.30hr) and 6 hours after heat-shock (bottom; after L3 div t=24.30hr) during the L3 division. V1, V2 and V3 lineages were followed over time; follow heat shock in L2, all anterior daughter cells behaved as seam cells in L3 (G) Spinning disk images of the seam cell syncytium in late L4 larvae that were control treated (top) or heat-shock exposed during the L2 (middle) or L3 (bottom) stage to induce *rnt-1 bro-1* expression. (H) Illustration of the endogenous *rnt-1* gene with introduced GFP-tag (Top). Time-lapse spinning disk microscopy of RNT-1::GFP and seam cell markers mCherry::PH and Cherry::H2B during L2 symmetric and asymmetric divisions. Images were processed using ImageJ software. Scale bars represent 10 µm.

By preventing differentiation, the RNT-1/BRO-1 complex could provide the mechanism that overrules the response to Wnt/β-catenin asymmetry during the symmetric L2 seam cell divisions. To examine this possibility, we set out to obtain further insight in the contribution of RNT-1 and BRO-1 in seam cell division and fate determination. First, we used heat-shock induced expression of RNT-1 and BRO-1 at times preceding the asymmetric divisions in L2 or L3. Following the L2 animals by time-lapse fluorescence microscopy (Fig. 4D and F) revealed that the anterior daughter cells of the normally asymmetric cell divisions failed to differentiate and fuse with hyp7 after induction of RNT-1/BRO-1. These cells continued to divide in L3 and behaved as normal seam daughter cells (Fig. 4D and F). Accordingly, quantification at the end of L4 larval development demonstrated that the seam cell numbers increased on average from 16 to 21 or 23 for animals heat-shock induced in L2 or L3, respectively (Fig. 4E and 4G). Thus, through temporal induction of RNT-1 and BRO-1, asymmetric seam cell divisions can be turned into symmetric divisions. In contrast, we did not observe additional seam cell divisions (Fig. 4D). Thus, ectopic expression highlights the contribution of RNT-1/BRO-1 in promoting the seam cell fate.

We wondered how soon after RNT-1/BRO-1 heat-shock induction the anterior daughter cells converted from a differentiation trajectory to a seam cell fate. Following normal asymmetric cell division, the two daughter cells differ immediately in cell cycle progression. The anterior daughter cells initiate the next cell cycle and undergo S phase prior to fusion with the hypodermis, while the posterior self-renewing seam cells pause in G0/G1 until the next molt (Hedgecock & White 1985). We previously visualized this difference in cell cycle progression with a CDK-activity sensor (Van Rijnberk et al. 2017). Nuclear export of this sensor, a DNA Helicase-GFP fusion protein, is induced by CDK-mediated phosphorylation. Consequently, after asymmetric division, S phase entry of the anterior daughter cell coincides with a reduced nuclear level of the CDK-sensor, while the quiescent posterior cell retains a high nuclear level (Fig. S1A-C). We induced expression of RNT-1 and BRO-1 just before the asymmetric L2 divisions, and followed the CDK-sensor in daughter cells after division. The nuclear GFP levels barely dropped in anterior daughter cells after RNT-1/BRO-1 induction (Fig. S1D). Thus, seam cells destined to divide asymmetrically switch to self-renewing seam cells within 90 minutes after heat-shock induced RNT-1/BRO-1 expression.

When combined with the time-lapse recordings of *rnt-1 bro-1* loss-of-function mutants, these data support the conclusion that RNT-1/BRO-1 promotes not only seam cell proliferation but also the seam cell fate. To study the normal expression of *rnt-1*, we used CRISPR/Cas9-assisted recombineering to insert gfp-coding sequences just before the translational stop codon in the endogenous gene (Fig. 4H, top). We followed RNT-1::GFP expression during larval development by time-lapse fluorescence microscopy. This revealed a high level of nuclear-localized RNT-1::GFP in interphase seam cells prior to the symmetric L2 divisions (Fig. 4H, Middle left). Interestingly, RNT-1::GFP subsequently disappeared during mitosis, and largely remained absent when the nuclei reformed in telophase (Fig. 4H, 25-45 minutes). This indicates active protein degradation and the possibility that RNT-1 is a substrate of the anaphase promoting complex/cyclosome (APC/C).

RNT-1::GFP did not reappear in the daughter cell nuclei prior to, or during, the L2 asymmetric divisions (Fig. 4H, Bottom rows). The APC/C becomes inactive prior to S phase entry, hence this is unlikely to be the only level of RNT-1 regulation. We considered post-transcriptional repression of *rnt-1* mRNA by microRNAs (miRNAs), which are important regulators of progression through L2, and transition to the L3 stage (Abbott et al. 2005; Li et al. 2005; Tsialikas et al. 2017). The *let-7* sister miRNAs, *miR-48, miR-84*, and *miR-241* would be candidates for *rnt-1* regulation, however, removal of a putative *let-7s* miRNA target site from the endogenous *rnt-1* 3’ untranslated region did not induce *rnt-1* gain of function (Fig. S3). Multiple levels of RNT-1 control are likely involved, and allow the reappearance of RNT-1 before the L3 stage division (Fig. S3). Importantly, the presence versus absence of nuclear RNT-1 prior to division distinguishes the symmetric division from the asymmetric division in L2 stage animals. The temporal control of RNT-1 expression in late L1 and L2 stage larvae, together with the L2 division phenotype in *rnt-1, bro-1, and unc-37* mutant larvae, indicate that the L2 symmetric seam cell divisions depend on transcriptional repression by RNT-1/BRO-1/UNC-37 in the mother seam cell.

### RNT-1/BRO-1 antagonize POP-1 at the level of anterior daughter cell differentiation

As a possible molecular mechanism, RNT-1/BRO-1 could overrule Wnt/β-catenin asymmetry during symmetric division by antagonizing POP-1 activity in anterior daughter cells. At a high nuclear level, POP-1 is thought to act as a transcriptional repressor, and to promote differentiation of anterior daughter cells (Kidd et al. 2005). By contrast, at a low nuclear level, POP-1 is expected to act as a transcriptional activator of Wnt-target genes, and to promote the stem cell-like fate of posterior daughter cells. However, RNAi of *pop-1* has been reported to strongly increase seam cell numbers, preventing the differentiation of anterior cells but not the stem-cell like seam cell fate (Gleason & Eisenmann 2010). This appears to indicate that only the repressor function of POP-1 is critical in the seam cell lineage. Complete absence of *pop-1* is lethal, however, and residual *pop-1* in the partial-loss-of-function RNAi animals could suffice for its activator function. To distinguish between these possibilities, we generated a conditional *pop-1* knockout allele. This was achieved by inserting *loxP*-recombination sites into the endogenous *pop-1* locus (Fig. 5A), and combining the homozygous loxed allele with seam-specific expression of the CRE recombinase (P*scm::CRE*; Ruijtenberg & van den Heuvel 2015).

**Figure 5.**
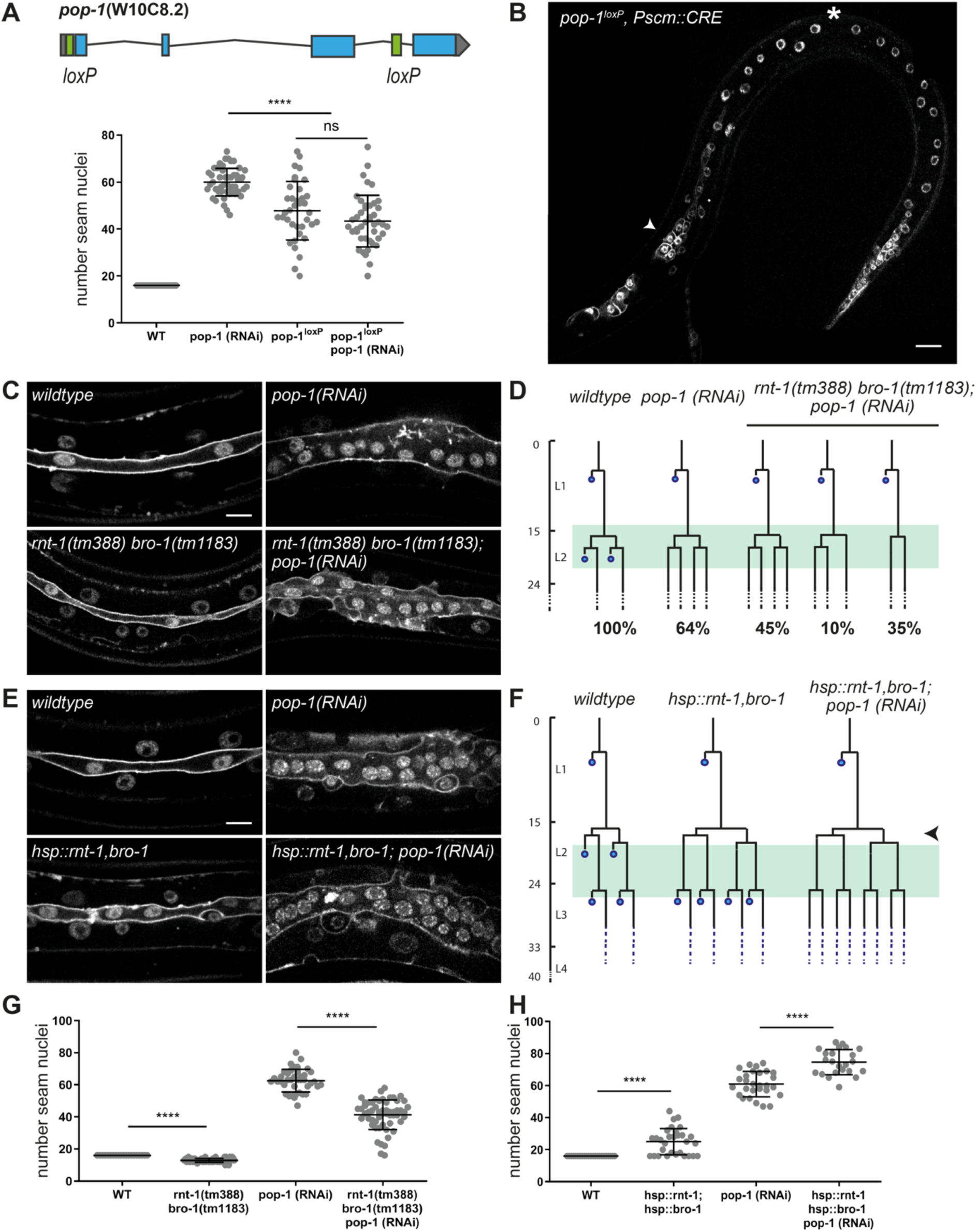
Possible antagonism between RNT-1/BRO-1 and POP-1 in controlling anterior seam cell differentiation. (A) Top: Gene map of the floxed endogenous *pop-1* allele. Bottom: Quantification of seam cell nuclei at the end of the L4 stage in *pop-1* RNAi, *pop-1*^loxP^ KO, and *pop-1*^loxP^ KO combined with *pop-1* feeding RNAi animals. (B) Spinning disk confocal microscopy image of a late L2 *pop-1*^loxP^, P*scm::CRE* animal. Seam markers are mCherry::PH and mCherry::H2B. Arrowhead points to extra seam cells, asterisk points to premature differentiation. (C) Spinning disk confocal microscopy images of late L4 wild-type or *rnt-1(tm388) bro-1(tm1183)* mutants, with and without *pop-1* RNAi. (D) Lineage analyses of L2 wild-type, *pop-1(RNAi)*, and *rnt-1(tm388) bro-1(tm1183), pop-1* RNAi larvae. Percentages refer to the fraction of seam cells displaying the phenotype. Green box marks time-window in which animals were observed. Spinning disk confocal microscopy image of heat-shock induced *rnt-1 bro-1* (late L4) with and without *pop-1* RNAi. (F) Lineage analyses of L2 heat-shock induced *rnt-1 bro-1* plus and minus *pop-1* RNAi. Time of heat-shock is marked by arrowhead. Green box indicates time-window during which animals were observed. (G) Quantification of seam cell nuclei at the end of L4 development for *rnt-1(tm388) bro-1(tm1183)* mutants plus and minus *pop-1* RNAi. (H) Quantification of seam cell nuclei at the end of L4 development for heat-shock-induced *rnt-1 bro-1* plus and minus *pop-1* RNAi. Images were processed with ImageJ software. Scale bars represent 10 µm. Error-bars represent mean ± SD.

Interestingly, the observed phenotype differed between the *pop-1* knockout and RNAi. RNAi knockdown increased the seam cell numbers to more than 60, as a result of anterior daughter cells failing to differentiate and adopting the seam fate (Fig. 5A). The seam-specific *pop-1* knockout also resulted in increased seam cell numbers, but to a lower extent. Closer examination showed that *pop-1*^*lox*^ knockout animals display a combination of anterior daughter cells adopting the seam fate, and abnormal differentiation of posterior seam cells (Fig. 5B and S4). Combined *pop-1* RNAi and lineage-specific knockout resembled the *pop-1* knockout alone (Fig. 5B). Thus, the exclusive failure in anterior cell differentiation following RNAi results from incomplete *pop-1* loss-of-function. The observations in the knockout agree with the paradigm that POP-1 exerts a dual role in seam daughter cells, promoting differentiation as a repressor, and the stem-cell fate as an activator. Because these functions are determined by the nuclear POP-1 levels, incomplete *pop-1* loss by RNAi likely removes the repressor but not activator function.

As RNT-1/BRO-1 suppresses seam cell differentiation, the complex could antagonize the differentiation-promoting *pop-1* repressor function. To test this possibility, we combined *pop-1* RNAi with heat-shock induced RNT-1/BRO-1 in L2 and L3 asymmetric divisions. This combination further increased the number of seam cells compared to either single condition (L2 heat-shock plus *pop-1* RNAi on average 75 seam cells, L3 heat-shock plus *pop-1* RNAi on average 95 cells. (Fig. 5E,F and 5H). This increase likely results from combining two incomplete conversions from asymmetric to symmetric seam cell division; ± 64% in L2 *pop-1*(*RNAi)* larvae (Fig. 5D, S5), versus one extra round of symmetric division (in L2 or L3) after heat-shock induction of RNT-1/BRO-1 (Fig. 4). Therefore, the enhanced phenotype of the *rnt-1 bro-1* (*gain of function*) *pop-1(RNAi)* combination compared to either single, does not indicate an order of gene functions or whether these genes act in a linear pathway. Nevertheless, these results confirm the antagonistic functions of the RNT-1/BRO-1 and POP-1 transcriptional regulators in anterior seam cell differentiation.

To further examine this antagonism, we combined *pop-1* RNAi with the *rnt-1 bro-1* double mutation. Seam nuclei counts at the end of L4 development revealed intermediate seam cell numbers for this combination (*rnt-1 bro-1* mutant 13 seam nuclei, *pop-1* RNAi 61 nuclei, *rnt-1 bro-1* combined with *pop-1* RNAi: 45 seam cell nuclei. Fig. 5C and 5G). Closer analysis of the seam cell lineages revealed that the ectopic differentiation of anterior daughter cells in *rnt-1 bro-1* mutants was completely suppressed by *pop-1* RNAi. The seam-cell proliferation defects of *rnt-1 bro-1* mutants, however, were not rescued by *pop-1* RNAi. This combination explains the intermediate seam cell numbers, and indicates that *pop-1* may act downstream of *rnt-1 bro-1*, specifically in differentiation control (Fig. 5D, G, and S5). Together, an antagonistic relation between RNT-1/BRO-1 and POP-1 is indicated by the overlap in phenotype between *rnt-1 bro-1* gain of function and *pop-1* loss of function, by the enhanced seam cell numbers that follow from combining *rnt-1 bro-1* gain of function and *pop-1* loss of function, and by the observed suppression of ectopic differentiation in *rnt-1 bro-1* mutants by *pop-1(RNAi)*. All these observations are consistent with the model that *rnt-1* and *bro-1* act upstream of *pop-1*, and inhibit differentiation by opposing the *pop-1* repressor function.

### RNT-1/BRO-1 antagonize POP-1 by negatively regulating its expression in L2 seam cells

The two most plausible scenarios by which the RNT-1/BRO-1 transcriptional repressor complex may negatively regulate POP-1 are either via transcriptional repression of *pop-1* itself, or via interfering with POP-1-mediated repression of Wnt target genes. In our initial experiments (Fig. 1A,B), we observed that POP-1 localizes asymmetrically during symmetric seam cell divisions. These experiments made use of *pop-1::gfp* expression from a multicopy integrated array, under the control of the *jmp#1* DNA fragment that turned out to be the *sys-1* promoter (Siegfried et al. 2004; LaBonty et al. 2014). As this transgene will not reflect normal POP-1 levels, we generated an *gfp*-tagged endogenous *pop-1* allele by CRISPR/Cas9-assisted recombineering (Fig. 6A). The homozygous *gfp::pop-1* strain was viable, although not fully healthy and occasionally missing a seam cell (Fig. S6). This indicates that while the tag is somewhat disruptive, GFP::POP-1 is largely functional. Hence, we used the GFP-tagged endogenous protein to determine POP-1 expression dynamics and the possibility of RNT-1/BRO-1-mediated suppression.

**Figure 6.**
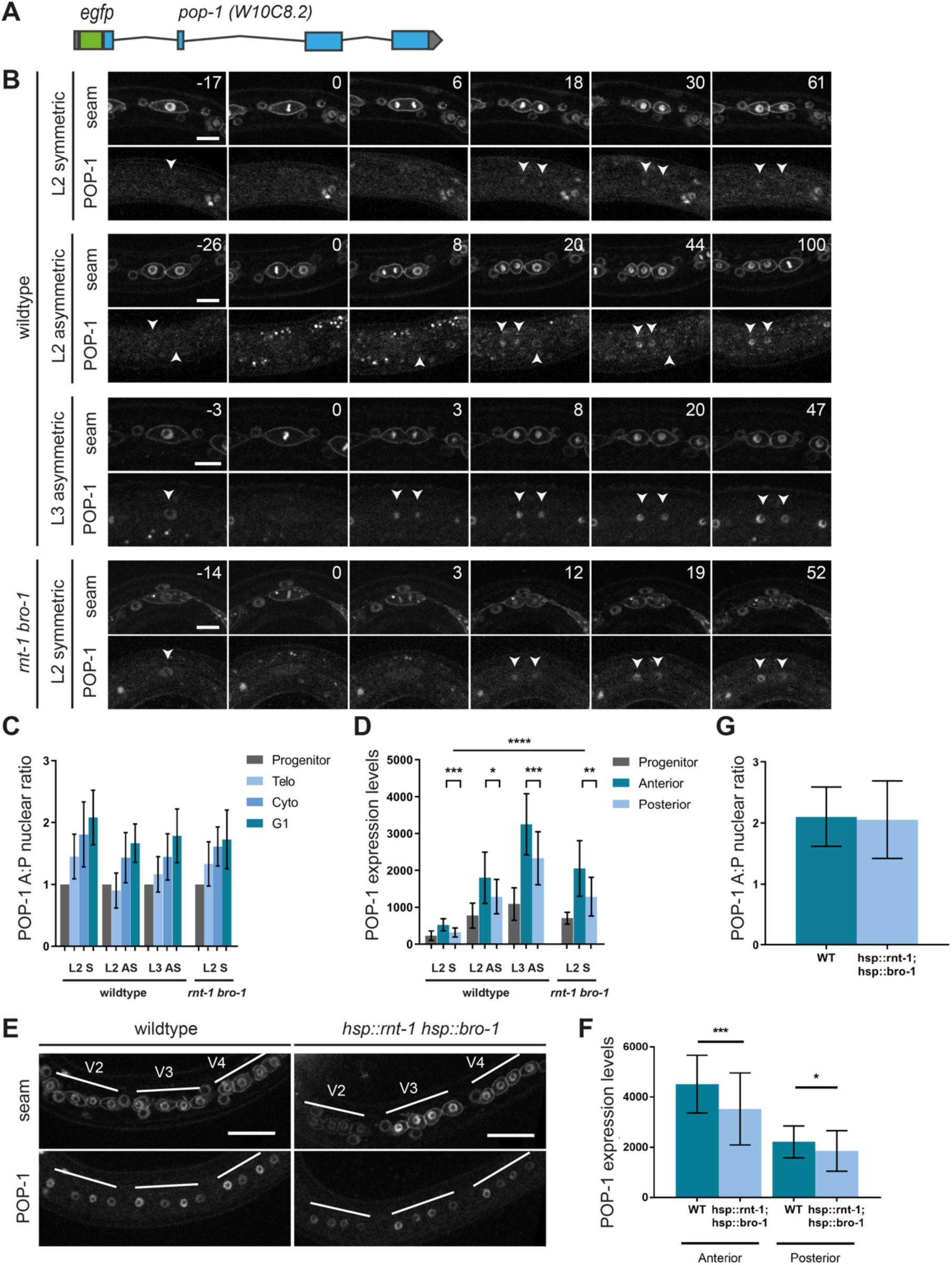
RNT-1/BRO-1 can overrule Wnt signaling by lowering *pop-1* expression levels. (A) Gene map of the tagged endogenous *gfp::pop-1* allele. (B) Time-lapse spinning disk confocal microscopy images of GFP::POP-1 and seam cell markers mCherry::PH and Cherry::H2B during L2 symmetric, L2 asymmetric, and L3 asymmetric divisions in the wild type, and L2 symmetric division in *rnt-1(tm388) bro-1(tm1183)* mutants. Arrowheads point to daughter cell nuclei. (C) Quantification of the A/P ratio of GFP::POP-1 in daughter cell nuclei of wildtype L2 and L3 divisions, and L2 symmetric division in *rnt-1(tm388) bro-1(tm1183)* mutants. (D) Relative expression levels of GFP::POP-1 during L2 and L3 divisions in the wild type, and L2 symmetric division in *rnt-1(tm388) bro-1(tm1183)* mutants. (E) Spinning disk confocal microscopy images of normal control and heat-shock induced RNT-1/BRO-1 late L2 animals. Seam markers are mCherry::PH and Cherry::H2B. (F) Relative GFP::POP-1 expression levels in control animals and heat-shock induced RNT-1/BRO-1 animals. (G) GFP::POP-1 nuclear A/P ratio in control animals and heat-shock induced RNT-1/BRO-1 animals. Note that the images in D and F were taken with different settings, therefore the relative levels (Y-axis) is different between these experiments. Images were processed using ImageJ software. Scale bars represent 10 µm (B) and 20 µm (E). Error-bars represent mean ± SD.

Similar to our observations of transgene-expressed *pop-1* (Fig. 2A,B), spinning disk confocal time-lapse microscopy of endogenous GFP::POP-1 showed asymmetric enrichment in anterior daughter cell nuclei formed during the symmetric L2 or asymmetric L2 and L3 divisions (Fig. 6B,C). Importantly, however, the POP-1 expression levels differed substantially between L2 and L3 seam cells: POP-1 levels were lowest during the symmetric division, and subsequently increased during L2 asymmetric and L3 asymmetric divisions (Fig. 6D; levels quantified at the time of cytokinesis). These observations suggest the possibility that RNT-1/BRO-1 overrule Wnt/β-catenin asymmetry in L2, by reducing *pop-1* expression.

To test this possibility, we compared GFP::POP-1 levels in wild-type and *rnt-1(tm388) bro-1(1183)* mutant animals. Notably, POP-1 expression levels were significantly higher in the mutant (Fig. 6B, bottom). In fact, during the first L2 seam cell division in the *rnt-1 bro-1* double mutant, GFP::POP-1 levels were similar to those of the asymmetric L2 division in the wild type (Fig. 6D). As nuclear POP-1 levels determine its activity as a transcriptional repressor, this finding may well explain that 31% of *rnt-1 bro-1* mutants skip the symmetric L2 division and show ectopic epidermal differentiation (Fig. 4A). To further test whether RNT-1/BRO-1 induces *pop-1* downregulation, we used heat-shock induced RNT-1 BRO-1 expression in the endogenous *gfp::pop-1* animals. This resulted in significantly reduced GFP::POP-1 levels in the daughter cells of the L2 asymmetric seam cell division (Fig. 6E-F). As expected, GFP::POP-1 still showed an asymmetric distribution between anterior and posterior daughter cells (Fig. 6G). We conclude that the RNT-1/BRO-1 transcriptional repressor is likely to reduce the expression of POP-1 below the threshold level needed for POP-1 repressor function, and thereby induces symmetric seam cell division. To test direct regulation, we altered two candidate RNT-1/BRO-1 binding sites in the *pop-1* promoter by CRISPR/Cas9-assisted recombineering (Fig. S7). This *pop-1* promoter mutation did not result in a *rnt-1 bro-1* phenotype, indicating that RNT-1/BRO-1 do not act (solely) through these elements.

## DISCUSSION

In this study, we examined the fundamental difference between asymmetric and symmetric seam cell divisions, and the mechanisms that control the switch between these division modes. In contrast to RNAi, lineage-specific *pop-1* knockout revealed the dual functions of POP-1 in the seam lineage. Only part of the *pop-1* knockout seam cells showed ectopic epidermal differentiation, while feeding RNAi resulted exclusively in failure to undergo differentiation of anterior daughter cells. The combined observations indicate that the transcriptional activator function of POP-1 is less critical than its repressor function, and requires a limited amount of POP-1. As removal of the repressor function is sufficient to convert an asymmetric seam cell division into a symmetric division, the presence or absence of POP-1-mediated transcriptional repression appears to be the fundamental difference between asymmetric and proliferative seam cell divisions.

Based on expression of a broadly used reporter transgene, we confirmed the earlier observation by us and others (Wildwater et al. 2011; Hughes et al. 2013; Harandi & Ambros 2014) that POP-1 levels differ between anterior and posterior daughter nuclei of symmetric divisions. Examination of GFP-tagged endogenous POP-1 confirmed this asymmetric localization. To our surprise, however, this also revealed a temporary decrease in *pop-1* expression prior to the L2 symmetric divisions, to a level substantially below that of nuclear POP-1 in self-renewing daughter cells of asymmetric seam cell divisions. A general reduction in POP-1 expression provides a simple mechanism to bypass the POP-1 repressor function during symmetric seam cell division.

### RNT-1/BRO-1 modulate Wnt signaling by negatively regulating *pop-1* gene expression

We identified the RNT-1/BRO-1 transcriptional repressor complex as a negative regulator of *pop-1* gene expression. Induced expression of RNT-1/BRO-1 resulted in symmetric cell division and significantly reduced GFP::POP-1 levels in seam cells. Conversely, loss of function of *rnt-1* and *bro-1* increased POP-1 expression in early L2 stage seam cells to a level normally present during the L2 asymmetric divisions. While supporting that RNT-1/BRO-1 negatively regulates POP-1 expression, these data do not reveal whether this regulation is direct. Supporting direct transcriptional regulation of *pop-1* by RNT-1/BRO-1, ChIP-sequencing results from the modERN consortium demonstrate RNT-1 association with the *pop-1* promoter in L1 larvae (Kudron et al. 2018). The *pop-1* promoter contains two Runx binding sites (5’-HGHGGK-3’; Van Der Deen et al., 2012) in this region. However, mutating these sites in the endogenous *pop-1* promoter did not result in a *rnt-1 bro-1* mutant phenotype (Fig. S7). It is possible that additional RNT-1/BRO-1 binding sites are present and sufficient for *pop-1* regulation. Alternatively, RNT-1/BRO-1 could downregulate *pop-1* indirectly, or contribute additional mechanisms to induce the L2 seam cell division program.

In L2, the presence versus absence of RNT-1 corresponds to POP-1 levels and symmetric versus asymmetric division. However, this is not true for other developmental stages. Tagged endogenous RNT-1 was highly expressed before the asymmetric seam cell divisions in L1 and L3 (Fig. S3), in line with observations with a transgenic reporter (Kagoshima et al. 2005). Similarly, analyses of reporter transgenes have indicated that BRO-1 is expressed through all larval stages (Kagoshima et al. 2007a; Xia et al. 2007). Interestingly, in males, the V6 seam cell undergoes an extra symmetric division during the L3 stage. This division and others in the male-specific V6 and T seam cell lineages are frequently skipped in *rnt-1* (also known as: male abnormal-2 *mab-2*) and *bro-1* mutants (Kagoshima et al. 2005, 2007a; Nimmo et al. 2005). It is possible that RNT-1/BRO-1 are more broadly expressed as an ancestral mechanism to induce symmetric seam cell divisions. In *C. elegans* this function is used only during L2 and male tail development, hence mechanisms need to be in place to prevent POP-1 repression at other stages. Studies of mammalian Runx proteins revealed extensive regulation by post-translational modifications that facilitate interaction with transcriptional activators or co-repressors and dictate Runx function (Reviewed in Blyth et al., 2005; Chuang et al., 2013). Similarly, the *C. elegans* RNT-1/BRO-1 repressor activity may be temporarily induced in L2 seam cells, or the response to RNT-1/BRO-1 activity could depend on other factors, such as the heterochronic pathway.

### Heterochronic genes may create a window of opportunity for *pop-1* repression

The temporal restriction of *pop-1* downregulation to the L2 stage seam cells suggests involvement of the heterochronic pathway. This pathway includes a series of successively expressed transcription factors, RNA-binding proteins and miRNAs that provide temporal identity during larval development (Rougvie 2005; Moss 2007). L1 development is determined by the LIN-14 transcription factor, which has been suggested to prevent symmetric seam cell division and reduce POP-1 dependence (Harandi & Ambros 2014). L2 development is defined by expression of the RNA-binding protein LIN-28 and downstream transcription factor HBL-1 (Abrahante et al. 2003; Lin et al. 2003; Abbott et al. 2005). These factors have also been shown to genetically interact with the Wnt/β-catenin asymmetry pathway in seam cells (Harandi & Ambros 2014). As a consequence, seam cells in the L2 stage appear uniquely sensitive to POP-1 levels, which could allow transitions from asymmetric to symmetric cell division. It is currently unclear whether this heterochronic effect could by mediated by activation of RNT-1/BRO-1, or inhibition of *pop-1* in parallel. Interestingly, a feedback loop between mammalian LIN28 and TCF7A has been detected in breast cancer cells (Chen et al. 2015), pointing to a potentially conserved mechanism. We did not observe an effect of heat-shock induced expression of either LIN-28 or HBL-1 during L2 and L3 asymmetric seam cell divisions (Fig. S8). Interestingly though, we did observe a genetic interaction between *rnt-1* and *hbl-1*; both *rnt-1* and *hbl-1* loss of function reduce the number of seam cell divisions in L2, and the combination strongly repressed the *pop-1(RNAi)* phenotype. Whether this reflects functions in parallel or within a regulatory cascade will require lineaging of null mutant combinations, as an extra division of seam nuclei in L4 *hbl-1(ve18)* larvae obscures the L2 cell division defective phenotype.

### Differential regulation of seam cell fate and proliferation

We did not observe additional divisions of seam cells following RNT-1/BRO-1 induction. The *rnt-1 bro-1* double mutant phenotype, however, supports that these factors also contribute to seam cell proliferation in the L2 stage, and during male tail development (Kagoshima et al. 2005, 2007a; Nimmo et al. 2005; Xia et al. 2007). RNAi of *pop-1* suppressed the ectopic differentiation but not proliferation defects of *rnt-1 bro-1* mutants, which indicates that cell fate and proliferation involve different mechanisms. The control of proliferation by RNT-1/BRO-1 has been suggested to involve repression of the cell cycle inhibitory genes *cki-1*^Cip/Kip^, *fzr-1*^Cdh1^ and *lin-35*^Rb^ (Nimmo et al. 2005; Kagoshima et al. 2007a; Xia et al. 2007). Analogous to the regulation of *pop-1* expression, it remains unclear how repression of these genes by RNT-1/BRO-1 is controlled to allow extra rounds of division only in the L2 stage and during male tail development. Similar to cell fate, the heterochronic factor LIN-28 could sensitize seam cells in the L2 stage for extra cell division. Mammalian Lin28 is a stem cell factor which promotes pluripotency and cell proliferation (Viswanathan & Daley 2010). Cyclin A, cyclin B and Cdk4 have been identified as target mRNAs for Lin28, and Lin28-mediated enhanced translation may promote stem cell proliferation (Xu et al. 2009). Similarly, upregulation of positive cell cycle regulators in L2 could determine that seam cells go through an extra division in response to RNT-1/BRO-1 mediated repression of cell-cycle inhibitors.

### Conserved modulation of Wnt signaling by Runx proteins

In this study, we identified a novel interaction between two conserved stem cell regulators. We propose that by negatively regulating *pop-1* expression, RNT-1/BRO-1 modulates Wnt/β-catenin asymmetry pathway activity in seam daughter cells (Summarized in Fig. 7). Cross-regulation between Runx and TCF appears conserved in mammals, although different mechanisms are likely involved. Studies in mouse intestinal epithelial cells showed that Runx3 adapts Wnt signaling via physical binding to nuclear TCF4. The formation of a ternary β-catenin::TCF4::Runx3 complex prevented TCF4 from binding to DNA (Ito et al. 2008; Reviewed in Chuang et al. 2013). Conversely, a ternary complex composed of β-catenin::LEF1::Runx2 was found to inhibit Runx2 from binding to DNA in mouse osteoblast cells (Kahler & Westendorf 2003). Whether or not such physical interactions are used in *C. elegans*, these results indicate that cross-regulation between the Runx/CBFβ and Wnt/β-catenin stem-cell regulators are likely applied more broadly.

**Figure 7.**
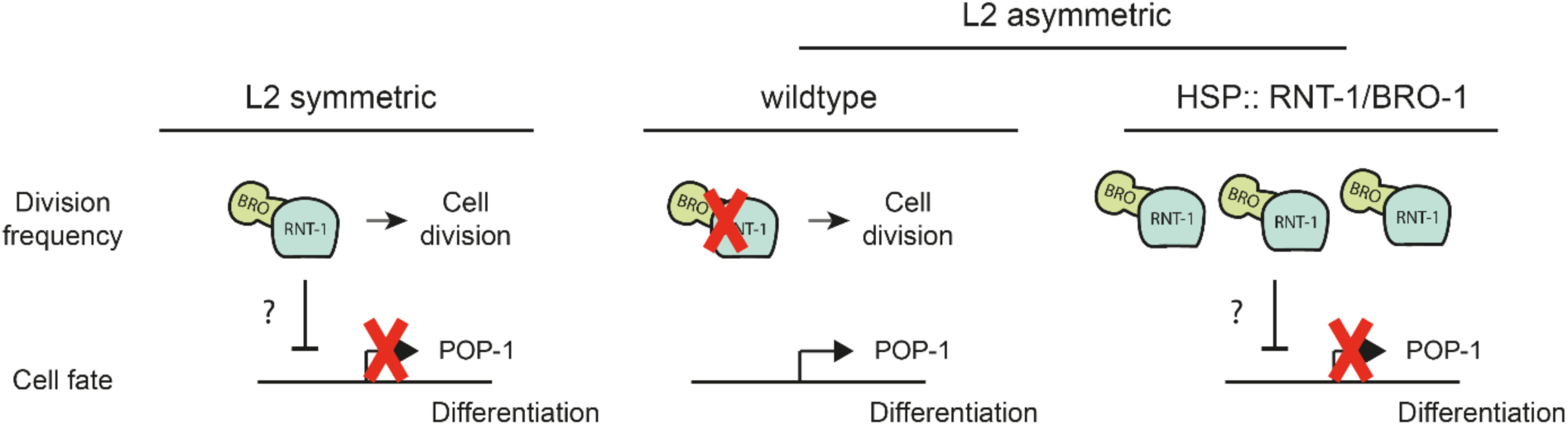
Model for RNT-1/BRO-1 mediated repression of POP-1. L2 seam cells maintain the asymmetric distribution of POP-1 during symmetric divisions. However, the RNT-1/BRO-1 repressor reduces the overall POP-1 expression level, and thereby nuclear POP-1 in the anterior daughter cell remains below the level needed for transcriptional repression and differentiation induction (left panel). Preceding the L2 asymmetric division in the wild-type, RNT-1 is degraded and POP-1 expression no longer repressed, allowing asymmetric cell division (middle). Increased expression levels of RNT-1/BRO-1 convert an asymmetric division into a symmetric division by reducing POP-1 expression levels (right panel).

## MATERIALS AND METHODS

### Nematode strains

Wild-type *Caenorhabditis elegans* strain N2 and the derivatives listed in Table 1 were used in this study. All strains were maintained at 20 °C as previously described (Brenner 1974) unless stated otherwise. Animals were grown on plates containing nematode growth medium seeded with OP50 *Escherichia coli* bacteria.

**Table 1.** Strains used in this study.

| Strain | Genotype |
| --- | --- |
| SV1009 | *hels63*[Pwrt-2::gfp::ph; Pwrt-2::gfp::h2b; Plin-48::mcherry] V |
| SV1984 | *hels218*[Pwrt-2::mcherry::ph, Pwrt-2::mcherry::h2b, Plin-48::gfp] IV |
| SV1986 | *hels218*[Pwrt-2::mcherry::ph, Pwrt-2::mcherry::h2b, Plin-48::gfp] IV; *qls74*[Psys-1::pop-1::gfp + *unc-119(+)*] *him-5*(e1490) V |
| JK3437 | *qls74*[Psys-1::pop-1::gfp + *unc-119(+)*] *him-5*(e1490) V; |
| HS1486 | *unc-76*(e911) V; *osls13*[*Papr-1::apr-1::venus*; *unc-76 (+)*] |
| SV1615 | *hels188* [Pwrt-2::mcherry::ph; Pwrt-2::mcherry::h2b; Plin-48::gfp; Phsp16.48::cki-1::gfp] |
| SV1694 | *unc-119* (*ed3*) III ; *heSi193* [Pmcm-4::CDK sensor::gfp::tbb-2 UTR + *Cbr-unc119(+)*] II ; *dpy-20* (*e1362*) ; [Pwrt-2::mcherry::ph Pwrt-2::mcherry::h2b *dpy-20(+)*] |
| SV2112 | *pop-1*(*he334*[*pop-1<sup>loxP</sup>*]) I* |
| SV2113 | *pop-1*(*he334*[*pop-1<sup>loxP</sup>*]) I; *hels218*[Pwrt-2::mcherry::ph, Pwrt-2::mcherry::h2b, Plin-48::gfp] IV; *heSi175*(Pscm::cre) X |
| SV2114 | *pop-1*(*he335*[*egfp::loxP::pop-1*]) I |
| SV2115 | *pop-1*(*he335*[*egfp::loxP::pop-1*]) I; *hels218*[Pwrt-2::mcherry::ph, Pwrt-2::mcherry::h2b, Plin-48::gfp] IV |
| SV2000 | *hels63*[Pwrt-2::gfp::ph; Pwrt-2::gfp::h2b; Plin-48::mcherry] V ; *heEx609*[Phsp16.2::rnt-1, Phsp16.2::bro-1, Pmyo-2::tdTomato] |
| SV2002 | *rnt-1*(*he305*[*rnt-1::egfp::3xflag::loxP*]) I |
| SV2003 | *rnt-1*(*he305*[*rnt-1::egfp::3xflag::loxP*]) I; *hels218*[Pwrt-2::mcherry::ph; Pwrt-2::mcherry::h2b; Plin-48::gfp] IV |
| YK138 | *rnt-1*(*tm388*) *bro-1*(*tm1183*) I |
| SV2126 | *rnt-1*(*tm388*) *bro-1*(*tm1183*) I; *hels63*[Pwrt-2::gfp::ph; Pwrt-2::gfp::h2b; Plin-48::mcherry] V |
| AW811 | *rnt-1*(*tm388*) I; *hels63*[Pwrt-2::gfp::ph; Pwrt-2::gfp::h2b; Plin-48::mcherry] V |
| SV2148 | *heSi193*[Pmcm-4::cdk-sensor] II; *hels218*[Pwrt-2::mcherry::ph, Pwrt-2::mcherry::h2b, Plin-48::gfp] IV; *heEx609*[Phsp16.2::rnt-1, Phsp16.2::bro-1, Pmyo-2::tdTomato] |
| SV2150 | *pop-1*(*he335*[*egfp::loxP::pop-1*]) I; *hels218*[Pwrt-2::mcherry::ph, Pwrt-2::mcherry::h2b, Plin-48::gfp] IV; *heEx609*[Phsp16.2::rnt-1, Phsp16.2::bro-1, Pmyo-2::tdTomato] |
| SV2230 | *pop-1*(*he335*[*egfp::loxP::pop-1*] *rnt-1*(*tm388*) *bro-1*(*tm1183*)) I; *hels218*[Pwrt-2::mcherry::ph, Pwrt-2::mcherry::h2b, Plin-48::gfp] IV |
| SV2231 | *pop-1*(*he373*[*pop-1<sup>ΔRunx</sup>*] I**); *hels63*[Pwrt-2::gfp::ph; Pwrt-2::gfp::h2b; Plin-48::mcherry] V |

\*loxP sites are present upstream of the *pop-1* ATG and in intron 5
\*\*promoter sites were altered between 143 and 164bp upstream of the *pop-1* ATG start codon (See Fig. S5)

### Molecular cloning

All molecular cloning was designed in ApE (A plasmid Editor; M. Wayne Davis). Repair templates and DNA fragments for cloning were generated by PCR amplification with either High Fidelity Hot Start KOD DNA Polymerase (Novagen) or Phusion Hot Start DNA Polymerase (Finnzymes), using either purified *C. elegans* genomic DNA or pre-existing vectors as template. A list of cloning primers can be found in Supplementary Table 1. PCR fragments were purified from gels (Qiagen), their concentration measured using a BioPhotometer D30 (Eppendorf) and then ligated into pCGSI by Gibson assembly (New England Biolabs) or pJJR82 (SEC cassette plus *egfp*; Dickinson et al. 2015). gRNA vectors were generated by annealing of antisense oligonucleotide pairs and subsequent ligation into pBbsI-linearized pJJR50 by T4 ligase (New England Biolabs). All DNA vectors used for genome editing were transformed into DH5*α* competent cells and subsequently purified by midiprep (Qiagen).

### CRISPR/Cas9 mediated genome editing

Knock-in strains were generated using Cas9 endonuclease-induced homologous recombination following standard methods (Dickinson et al. 2013). Repair templates were generated by inserting 500 bp homology arm PCR products into destination vectors containing *egfp* and a self-excising selection cassette using Gibson assembly (New England Biolabs) or SapTrap assembly (Dickinson et al. 2015; Schwartz & Jorgensen 2016). Destination vectors used in this study were pJJR82 (C-terminal *rnt-1*) and pMLS257 (N-terminal *pop-1*). The C-terminal *rnt-1* repair template contains a nine amino acid flexible linker between the coding sequence and the *egfp* tag. N-terminal *pop-1* repair template contains a ten amino acid flexible linker between the coding sequence and the *egfp* tag. Injection of *C. elegans* adults in the germline was performed using an inverted microinjection microscope setup. Injection mixes with a total volume of 50 μl were prepared in milliQ H_2_O and contained a combination of 30-50 ng/μl *Peft-3::cas9* (46168; Addgene; Friedland et al., 2013), 50-100 ng/μl *Pu6::sgRNA* with sequences targeted against *pop-1* or *rnt-1*, 50 ng/μl repair vector, and 2.5 ng/μl *Pmyo-2::tdTomato* as a co-injection marker. Injected animals were transferred to new NGM-OP50 plates (3 animals per plate) and allowed to lay eggs for 2-3 days at 25 °C. On day 3, 500 μl of filter sterilized hygromycin solution (5 mg/ml in water) was added to the plates and allowed to dry in. Plates were subsequently moved back to 25 °C. On day 7, plates were screened for surviving F1 animals that showed a Rol phenotype and lacked the co-injection marker. These candidate knock-in animals were singled to new NGM-OP50 plates without hygromycin. On day 10, plates with homozygous Rol progeny (C-terminal *rnt-1*) and heterozygous Rol animals (N-terminal *pop-1*) were selected. Of those, 6 L1 animals were transferred to new plates, and exposed to heat-shock at 34 °C for 4 hours for cassette excision. Subsequent genome editing events were assessed by microscopic analysis and PCR amplification using primers targeting the inserted *gfp* sequence and a genomic region outside the homology arms. PCR-confirmed edited genomic loci were further validated by DNA sequencing (Macrogen Europe).

*loxP* sites were integrated in the endogenous locus of *pop-1* via co-conversion in a *pha-1(e2123*ts) background. Injection mixes contained a combination of 30-50 ng/μl *Peft-3::cas9* (46168; Addgene; Friedland et al., 2013), 50-100 ng/μl *Pu6::sgRNA* with sequences targeted against *pop-1*, 50 ng/μl of PAGE-purified *pop-1* repair oligo (Integrated DNA technologies), 50 ng/μl PAGE-purified *pha-1* repair oligo (Integrated DNA technologies), 60 ng/μl pJW1285 (61252; Addgene; Ward, 2015), and 2.5 ng/μl Pmyo-2::tdTomato as a co-injection marker. Animals were grown for 3-5 days at either 20 °C or 25 °C after injection, and transgenic progeny was selected based on either expression of tdTomato in the pharynx or survival at the non-permissive temperature (25 °C). Subsequent assessment of genome editing events was performed by PCR amplification using primers targeting the inserted *loxP* sequence and genomic sequences outside the homology arms. PCR-confirmed edited genomic loci were further validated by DNA sequencing (Macrogen Europe).

### Generation of extrachromosomal arrays

Extrachromosomal arrays were generated for *hsp::rnt-1 (pAW261), hsp::bro-1 (pAW266), hsp::cki-1::gfp*, and P*wrt-2::mCherry::PH*, P*wrt-2::mCherry::H2B*. For heat-shock induced RNT-1/BRO-1 expression, the injection mix contained a combination of 30 ng/μl P*hsp16*.*2::rnt-1*, 30 ng/μl P*hsp16*.*2::bro-1*, 2.5 ng/μl P*myo-2::tdTomato* and 5 ng/μl *λ*-DNA. For the seam markers, the injection mix contained 50 ng/μl P*wrt-2::mcherry::ph*, 50 ng/μl P*wrt-2::mcherry::h2b*, 10 ug/μl P*lin-48::GFP* and 5 ng/μl *λ*-DNA. For CKI-1 induction, the injection mix contained a combination of 20 ng/μl P*hsp16*.*48::cki-1::gfp*, 2.5 ug/μl P*myo-2::tdTomato* and 5 ng/μl *λ*-DNA. Animals were grown for 3-5 days at 20 °C, and transgenic progeny was selected based on pharyngeal expression of tdTomato (*myo-2*) or tail expression of GFP (*lin-48*). Strains were maintained as extrachromosomal lines by transferring tdTomato or GFP positive animals.

Integration by γ-irradiation was performed for extrachromosomal arrays containing *hsp16*.*48::cki-1::gfp* and the combined P*wrt-2::mcherry::h2b* and P*wrt-2::mcherry::ph* markers.

### Staging

Animals were synchronized using a wash-off protocol. 20 gravid adults were transferred to a new NGM-OP50 plate and allowed to lay eggs for a minimum of 20 hours. Animals were washed off the plates using M9-0,1%Tween, and embryos were allowed to hatch for a period of 1 hour. The newly hatched larvae were collected onto a fresh NMG-OP50 plate and incubated at 20 °C for 4.5 hours (L1), 15.5 hours (L2 symmetric), 17.5 hours (L2 asymmetric), 24 hours (L3) or 43 hours (late L4 counting).

### RNA-mediated interference (RNAi)

A combination of L1 soaking and feeding RNAi was used to knock-down *pop-1*. Gravid adults were bleached using hypochlorite treatment, and embryos were allowed to hatch for 20 hours in RNAi soaking buffer (0.05% gelatin, 5.5 mM KH_2_PO_4_, 2.1 mM NaCl, 4.7 mM NH_4_Cl, 3 mM spermidine) containing 1 ug dsRNA. Hatched larvae were then placed on 5x concentrated RNAi feeding plates at 20°C. Both the RNAi feeding plates and the dsRNA were derived from Vidal library clone GHR-11053 for *pop-1*. dsRNA was synthesized using the Megascript High Yield Transcription T7 Kit (Thermofisher Scientific).

### Heat-shock induction

For heat-shock induced gene expression, animals were synchronized using a wash-off protocol (see ‘Staging’) and grown at 20 °C. Heat-shock was performed in a 32 °C water bath for 30 min (CKI-1) or 60 min (RNT-1/BRO-1). After heat-shock, the plates were placed on ice-water for 10 minutes and either used directly for microscopy or placed back at 20 °C for later analysis.

### Microscopy

Time-lapse movies of seam cell divisions in immobilized, living animals were recorded at room-temperature at 2-minute intervals for 2-5 hours using a Nikon Eclipse Ti-U spinning disk microscope with a 63X objective. Larvae were immobilized in 1 mM tetramisole (Sigma-Aldrich) in M9 buffer, and mounted on 5-7% agarose pads (7% for L1 stage animals, 5% for L2-L3 stage animals. Agarose was prepared in milliQ water). The coverslips were sealed with immersion oil (Zeiss Immersol 518N oil) to prevent liquid evaporation. Laser power (both 488 and 563) ranged between 6-10% with exposure times below 400 ms for long-term imaging. 2×2 binning was performed to reduce phototoxicity. Image analysis was performed with FIJI software. Quantification of endogenous expression levels was corrected for background levels inside the worm.

## Acknowledgements

The authors thank members of the van den Heuvel, Boxem and Woollard labs for help and discussion. We thank Eugene Katrukah (A. Akhmanova lab) for help and expertise with analyzing fluorescence intensities. Some strains were provided by the CGC, which is funded by NIH Office of Research Infrastructure Programs (P40 OD010440).

## References

Abbott, A.L., Alvarez-Saavedra, E., Miska, E.A., Lau, N.C., Bartel, D.P., Horvitz, H.R. & Ambros, V. (2005) The let-7 MicroRNA family members mir-48, mir-84, and mir-241 function together to regulate developmental timing in Caenorhabditis elegans. Developmental Cell 9: 403–414.

Abrahante, J.E., Daul, A.L., Li, M., Volk, M.L., Tennessen, J.M., Miller, E.A., Rougvie, A.E., Hall, J. & Se, C.S. (2003) The Caenorhabditis elegans hunchback-like Gene lin-57 / hbl-1 Controls Developmental Time and Is Regulated by MicroRNAs. 4: 625–637.

Ambros, V. (1989) A hierarchy of regulatory genes controls a larva-to-adult developmental switch in C. elegans. Cell 57: 49–57.

Ambros, V.R. & Horvitz, H.R. (1984) Heterochronic Mutants of the Nematode Caenorhabditis elegans Authors (s): Victor Ambros and H. Robert Horvitz Source: Science, New Series, Vol. 226, No. 4673 (Oct. 26, 1984), pp. 409-416 Published by: American Association for the Advancemen. Science 226: 409–416.

Baldwin, A.T. & Phillips, B.T. (2014) The tumor suppressor APC differentially regulates multiple βcatenins through the function of axin and CKIα during C. elegans asymmetric stem cell divisions. Journal of cell science 127: 2771–81.

Banerjee, D., Chen, X., Lin, S.Y. & Slack, F.J. (2010) kin-19/casein kinase Ia has dual functions in regulating asymmetric division and terminal differentiation in C. elegans epidermal stem cells. Cell Cycle 9: 4748–4765.

Blyth, K., Cameron, E.R. & Neil, J.C. (2005) The RUNX genes: Gain or loss of function in cancer. Nature Reviews Cancer 5: 376–387.

Brenner, S. (1974) The genetics of Caenorhabditis elegans. Genetics 77: 71–94.

Calvo, D., Victor, M., Gay, F., Sui, G., Luke, M.P.S., Dufourcq, P., Wen, G., Maduro, M., Rothman, J. & Shi, Y. (2002) A POP-1 repressor complex restricts inappropriate cell type-specific gene transcription during Caenorhabditis elegans embryogenesis. EMBO Journal 20: 7197–7208.

Chen, C., Cao, F., Bai, L., Liu, Y., Xie, J., Wang, W., Si, Q., Yang, J., Chang, A., Liu, D., Liu, D., Chuang, T.H., Xiang, R. & Luo, Y. (2015) IKKβ enforces a LIN28B/TCF7L2 positive feedback loop that promotes cancer cell stemness and metastasis. Cancer Research 75: 1725–1735.

Chuang, L.S.H., Ito, K. & Ito, Y. (2013) RUNX family: Regulation and diversification of roles through interacting proteins. International Journal of Cancer 132: 1260–1271.

Van Der Deen, M., Akech, J., Lapointe, D., Gupta, S., Young, D.W., Montecino, M.A., Galindo, M., Lian, J.B., Stein, J.L., Stein, G.S. & Van Wijnen, A.J. (2012) Genomic promoter occupancy of Runt-related transcription factor RUNX2 in osteosarcoma cells identifies genes involved in cell adhesion and motility. Journal of Biological Chemistry 287: 4503–4517.

Deltcheva, E. & Nimmo, R. (2017) RUNX transcription factors at the interface of stem cells and cancer. Biochemical Journal 474: 1755–1768.

Dickinson, D.J., Pani, A.M., Heppert, J.K., Higgins, C.D. & Goldstein, B. (2015) Streamlined genome engineering with a self-excising drug selection cassette. Genetics 200: 1035–1049.

Dickinson, D.J., Ward, J.D., Reiner, D.J. & Goldstein, B. (2013) Engineering the Caenorhabditis elegans genome using Cas9-triggered homologous recombination. Nature Methods 10: 1028–1034.

Friedland, A.E., Tzur, Y.B., Esvelt, K.M., Colaiácovo, M.P., Church, G.M. & Calarco, J.A. (2013) Heritable genome editing in C. elegans via a CRISPR-Cas9 system. Nature Methods 10: 741–743.

Gleason, J.E. & Eisenmann, D.M. (2010) Wnt signaling controls the stem cell-like asymmetric division of the epithelial seam cells during C. elegans larval development. Developmental Biology 348: 58–66.

Gorrepati, L., Thompson, K.W. & Eisenmann, D.M. (2013) C. elegans GATA factors EGL-18 and ELT-6 function downstream of Wnt signaling to maintain the progenitor fate during larval asymmetric divisions of the seam cells. Development (Cambridge, England) 140: 2093–102.

Harandi, O.F. & Ambros, V.R. (2014) Control of stem cell self-renewal and differentiation by the heterochronic genes and the cellular asymmetry machinery in Caenorhabditis elegans. Proceedings of the National Academy of Sciences 112: E287–E296.

Hedgecock, E.M. & White, J.G. (1985) Polyploid tissues in the nematode Caenorhabditis elegans. Developmental biology 107: 128–133.

Huang, S., Shetty, P., Robertson, S.M. & Lin, R. (2007) Binary cell fate specification during C. elegans embryogenesis driven by reiterated reciprocal asymmetry of TCF POP-1 and its coactivator beta-catenin SYS-1. Development 134: 2685–95.

Hughes, S., Brabin, C., Appleford, P.J. & Woollard, A. (2013) CEH-20/Pbx and UNC-62/Meis function upstream of rnt-1/Runx to regulate asymmetric divisions of the C. elegans stem-like seam cells. Biology open 2: 718–27.

Ito, K., Lim, A.C.B., Salto-Tellez, M., Motoda, L., Osato, M., Chuang, L.S.H., Lee, C.W.L., Voon, D.C.C., Koo, J.K.W., Wang, H., Fukamachi, H. & Ito, Y. (2008) RUNX3 Attenuates β-Catenin/T Cell Factors in Intestinal Tumorigenesis. Cancer Cell 14: 226–237.

Kagoshima, H., Nimmo, R., Saad, N., Tanaka, J., Miwa, Y., Mitani, S., Kohara, Y. & Woollard, A. (2007a) The C. elegans CBFbeta homologue BRO-1 interacts with the Runx factor, RNT-1, to promote stem cell proliferation and self-renewal. Development 134: 3905–3915.

Kagoshima, H., Nimmo, R., Saad, N., Tanaka, J., Miwa, Y., Mitani, S., Kohara, Y. & Woollard, A. (2007b) The C. elegans CBFbeta homologue BRO-1 interacts with the Runx factor, RNT-1, to promote stem cell proliferation and self-renewal. Development (Cambridge, England) 134: 3905–3915.

Kagoshima, H., Sawa, H., Mitani, S. & Bu, T.R. (2005) The C. elegans RUNX transcription factor RNT-1 / MAB-2 is required for asymmetrical cell division of the T blast cell. 287: 262–273.

Kahler, R.A. & Westendorf, J.J. (2003) Lymphoid enhancer factor-1 and β-catenin inhibit Runx2-dependent transcriptional activation of the osteocalcin promoter. Journal of Biological Chemistry 278: 11937–11944.

Kidd, A.R., Miskowski, J.A., Siegfried, K.R., Sawa, H. & Kimble, J. (2005) A β-catenin identified by functional rather than sequence criteria and its role in Wnt/MAPK signaling. Cell 121: 761–772.

Knoblich, J.A. (2010) Asymmetric cell division: recent developments and their implications for tumour biology. Nature Reviews Molecular Cell Biology 11: 849–860.

Kudron, M.M., Victorsen, A., Gevirtzman, L., Hillier, L.W., Fisher, W.W., Vafeados, D., Kirkey, M., Hammonds, A.S., Gersch, J., Ammouri, H., Wall, M.L., Moran, J., Steffen, D., Szynkarek, M., Seabrook-Sturgis, S., Jameel, N., Kadaba, M., Patton, J., Terrell, R., Corson, M., Durham, T.J., Park, S., Samanta, S., Han, M., Xu, J., Yan, K.K., Celniker, S.E., White, K.P., Ma, L., Gerstein, M., Reinke, V. & Waterston, R.H. (2018) The modern resource: genome-wide binding profiles for hundreds of Drosophila and Caenorhabditis elegans transcription factors. Genetics 208: 937– 949.

LaBonty, M., Szmygiel, C., Byrnes, L.E., Hughes, S., Woollard, A. & Cram, E.J. (2014) CACN-1/Cactin plays a role in Wnt signaling in C. elegans. PloS one 9: e101945.

Li, M., Jones-Rhoades, M.W., Lau, N.C., Bartel, D.P. & Rougvie, A.E. (2005) Regulatory mutations of mir-48, a C. elegans let-7 family microRNA, cause developmental timing defects. Developmental Cell 9: 415–422.

Lin, R., Hill, R.J. & Priess, J.R. (1998) POP-1 and anterior-posterior fate decisions in C. elegans embryos. Cell 92: 229–239.

Lin, S.Y., Johnson, S.M., Abraham, M., Vella, M.C., Pasquinelli, A., Gamberi, C., Gottlieb, E. & Slack, F.J. (2003) The C. elegans hunchback homolog, hbl-1, controls temporal patterning and is a probable MicroRNA target. Developmental Cell 4: 639–650.

Mizumoto, K. & Sawa, H. (2007a) Cortical beta-catenin and APC regulate asymmetric nuclear beta-catenin localization during asymmetric cell division in C. elegans. Developmental cell 12: 287– 299.

Mizumoto, K. & Sawa, H. (2007b) Two betas or not two betas: regulation of asymmetric division by beta-catenin. Trends in Cell Biology 17: 465–473.

Moss, E.G. (2007) Heterochronic Genes and the Nature of Developmental Time. Current Biology 17: 425–434.

Moss, E.G., Lee, R.C. & Ambros, V. (1997) The cold shock domain protein LIN-28 controls developmental timing in C. elegans and is regulated by the lin-4 RNA. Cell 88: 637–646.

Nimmo, R., Antebi, A. & Woollard, A. (2005) mab-2 encodes RNT-1, a C. elegans Runx homologue essential for controlling cell proliferation in a stem cell-like developmental lineage. Development 132: 5043–5054.

Ren, H. & Zhang, H. (2010) Wnt signaling controls temporal identities of seam cells in Caenorhabditis elegans. Developmental Biology 345: 144–155.

Van Rijnberk, L.M., Van Der Horst, S.E.M., Van Den Heuvel, S. & Ruijtenberg, S. (2017) A dual transcriptional reporter and CDK-activity sensor marks cell cycle entry and progression in C. elegans. PLoS ONE 12: 1–14.

Rougvie, A.E. (2005) Intrinsic and extrinsic regulators of developmental timing: from miRNAs to nutritional cues. Development (Cambridge, England) 132: 3787–3798.

Ruijtenberg, S. & van den Heuvel, S. (2016) Coordinating cell proliferation and differentiation: Antagonism between cell cycle regulators and cell type-specific gene expression. Cell Cycle 15: 196–212.

Ruijtenberg, S. & Van Den Heuvel, S. (2015) G1/S Inhibitors and the SWI/SNF Complex Control Cell-Cycle Exit during Muscle Differentiation. Cell 162: 300–313.

Schwartz, M.L. & Jorgensen, E.M. (2016) SapTrap, a Toolkit for High-Throughput CRISPR / Cas9. 202: 1277–1288.

Shetty, P., Lo, M.C., Robertson, S.M. & Lin, R. (2005) C. elegans TCF protein, POP-1, converts from repressor to activator as a result of Wnt-induced lowering of nuclear levels. Developmental Biology 285: 584–592.

Siegfried, K.R., Kidd, A.R., Chesney, M. a. & Kimble, J. (2004) The sys-1 and sys-3 Genes Cooperate with Wnt Signaling to Establish the Proximal-Distal Axis of the Caenorhabditis elegans Gonad. Genetics 166: 171–186.

Slack, F.J., Basson, M., Liu, Z., Ambros, V., Horvitz, H.R. & Ruvkun, G. (2000) The lin-41 RBCC gene acts in the C. elegans heterochronic pathway between the let-7 regulatory RNA and the LIN-29 transcription factor. Molecular cell 5: 659–669.

Slack, F.J. & Ruvkun, G. (1997) Temporal pattern formation by heterochronic genes. Annual review of genetics 31: 611–34.

Sulston, J.E. & Horvitz, H.R. (1977) Post-embryonic cell lineages of the nematode, Caenorhabditis elegans. Developmental Biology 56: 110–156.

Sulston, J.E., Schierenberg, E., White, J.G. & Thomson, J.N. (1983) The embryonic cell lineage of the nematode Caenorhabditis elegans. Developmental Biology 100: 64–119.

Takeshita, H. & Sawa, H. (2005) Asymmetric cortical and nuclear localizations of WRM-1/??-catenin during asymmetric cell division in C. elegans. Genes and Development 19: 1743–1748.

Tsialikas, J., Romens, M.A., Abbott, A. & Moss, E.G. (2017) Stage-specific timing of the microRNA regulation of lin-28 by the heterochronic gene lin-14 in Caenorhabditis elegans. Genetics 205: 251–262.

Viswanathan, S.R. & Daley, G.Q. (2010) Lin28: A MicroRNA Regulator with a Macro Role. Cell 140: 445–449.

Vora, S. & Phillips, B.T. (2015) Centrosome-Associated Degradation Limits β-Catenin Inheritance by Daughter Cells after Asymmetric Division. Current biology: CB 25: 1005–16.

Ward, J.D. (2015) Rapid and precise engineering of the caenorhabditis elegans genome with lethal mutation co-conversion and inactivation of NHEJ repair. Genetics 199: 363–377.

Wildwater, M., Sander, N., de Vreede, G. & van den Heuvel, S. (2011) Cell shape and Wnt signaling redundantly control the division axis of C. elegans epithelial stem cells. Development 138: 4375–4385.

Xia, D., Zhang, Y., Huang, X., Sun, Y. & Zhang, H. (2007) The C. elegans CBFbeta homolog, BRO-1, regulates the proliferation, differentiation and specification of the stem cell-like seam cell lineages. Developmental biology 309: 259–272.

Xu, B., Zhang, K. & Huang, Y. (2009) Lin28 modulates cell growth and associates with a subset of cell cycle regulator mRNAs in mouse embryonic stem cells. RNA (New York, N.Y.) 15: 357–361.

