## Supplementary Information for "*C. elegans* Runx/CBFβ suppresses POP-1(TCF) to convert asymmetric to proliferative division of stem cell-like seam cells"

SUPPLEMENTARY INFORMATION VAN DER HORST ET AL., 2019

### Figure S1

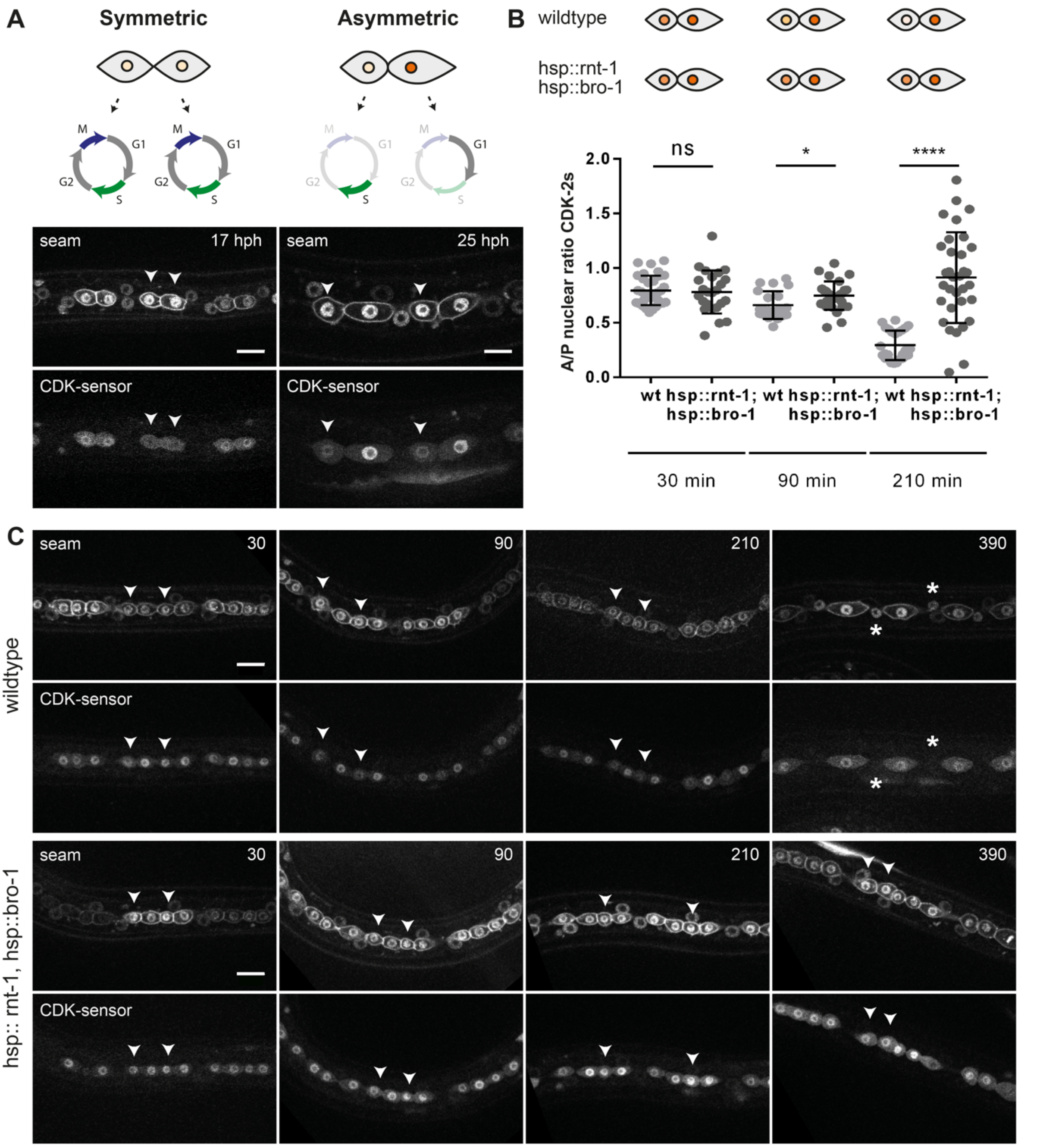


**Figure S1. Cell cycle progression as a read-out for daughter cell fate.** (A) Seam cells progress differently through the cell cycle after symmetric or asymmetric cell division (Top), as illustrated by spinning disk microscopy images of seam cells expressing the CDK sensor (Bottom). Left panels: seam daughter cells after symmetric L2 division, t=17 hours post hatching (hph). Right panels: daughter cells after asymmetric division in L3, t=25 hph. Note that the CDK sensor helps to distinguish between anterior fate (cell cycle reentry; nuclear export of GFP) and seam cell self-renewal (quiescence; nuclear retention of GFP). (B) A/P nuclear ratio of the CDK-2 sensor in control and heat-shock induced RNT-1/BRO-1 L2 seam cells measured at different timepoints after induction. Time indicates minutes after heat shock, which was stopped just prior to or during mitosis. (C) Spinning disk microscopy of control and heat-shock induced RNT-1/BRO-1 seam cells. Images taken at different time point after induction. Images were processed using ImageJ software. Scale bars represent 10 µm. Error-bars represent mean ± SD.

**Figure S2**

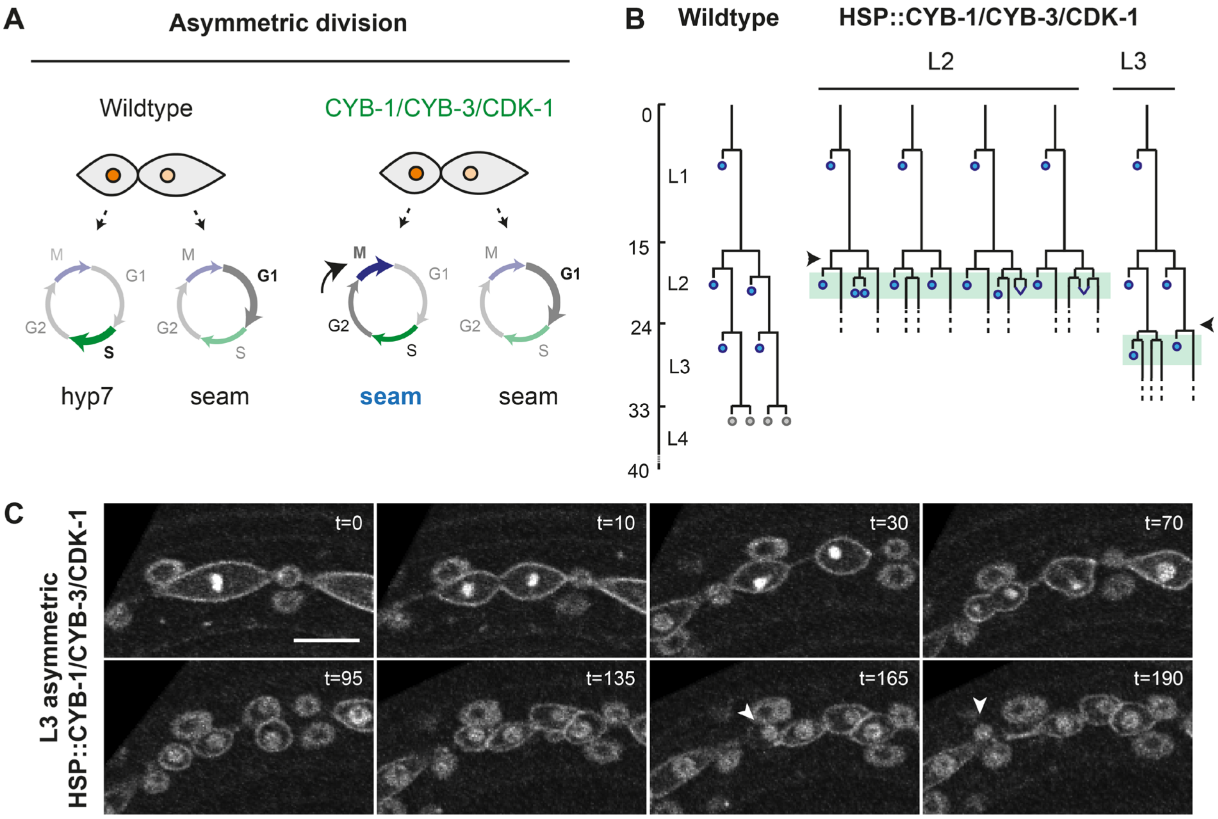


**Figure S2. Induced expression of CDK-1, CYB-1, and CYB-3 occasionally induced extra mitosis.** (A) Seam daughters of a wild-type asymmetric division progress differently after cytokinesis: anterior differentiating daughter cells undergo S-phase before fusing with the hyp7 epidermis. Posterior daughters pause in G1 until the next larval stage. Heat-shock induced expression of *cdk-1, cyb-1,* and *cyb-3* was used to force anterior daughter cells of asymmetric divisions to progress into mitosis after completing S-phase. We hypothesized that cell cycle progression of the anterior cells might overrule differentiation into hyp7 and trigger these cells to maintain a seam fate (blue). (B). Lineage analyses of L2 (middle) and L3 (right) animals with heat-shock induced CDK-1/CYB-1/CYB-3. Heat shock was given between the symmetric and asymmetric L2 division (middle; arrowhead) or prior to the L3 asymmetric division (right; arrowhead). Animals were followed using time-lapse microscopy during the hours after heat shock (green boxes). (C) Time-lapse spinning disk microscopy of L3 animals with heat-shock induced CDK-1/CYB-1/CYB-3. Time-lapse represents the L3 lineage presented in panel B. Both the anterior and posterior daughter cells of the L3 asymmetric division undergo an additional mitosis, of which the anterior daughter cell of the anterior division differentiates and fuses with the epidermis (arrowhead). Images were processed using ImageJ software. Scale bar represents 10 µm.

### Figure S3

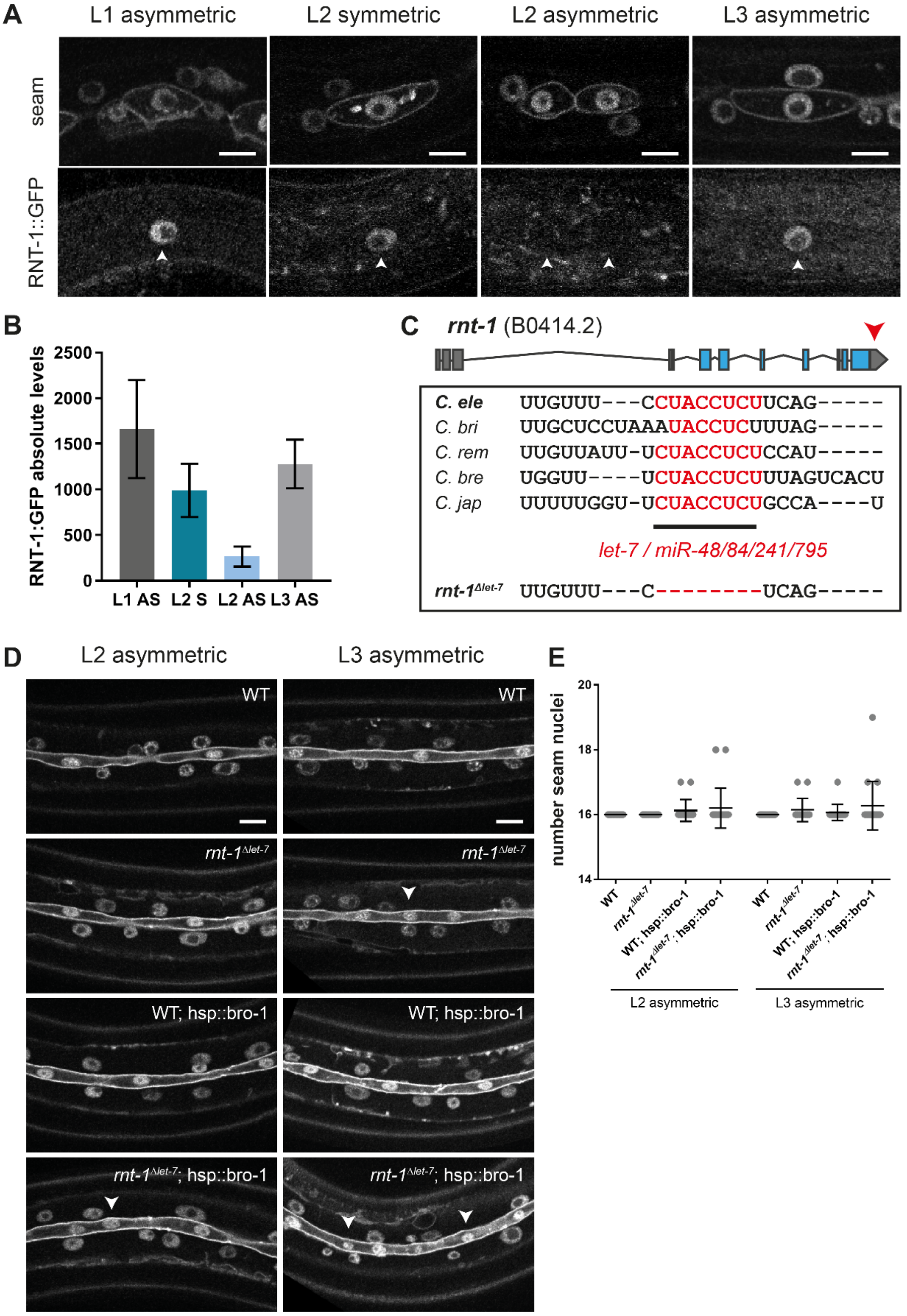

A
L1 asymmetric
L2 symmetric
L2 asymmetric
L3 asymmetric
seam
RNT-1::GFP
B
RNT-1:GFP absolute levels
2500
2000
1500
1000
500
0
L1 AS
L2 S
L2 AS
L3 AS
C
rnt-1 (B0414.2)
C. ele UUGUUU---CCUACCUCUUCAG-----
C. bri UUGCUCCUAAAUACCUCUUUAG-----
C. rem UUGUUAUU-UCUACCUCUCCAU-----
C. bre UGGUU----UCUACCUCUUUAGUCACU
C. jap UUUUUGGU-UCUACCUCUGCCA----U
let-7 / miR-48/84/241/795
rnt-1Δlet-7 UUGUUU---C--------UCAG-----
D
L2 asymmetric
L3 asymmetric
WT
rnt-1Δlet-7
WT; hsp::bro-1
rnt-1Δlet-7; hsp::bro-1
E
number seam nuclei
20
18
16
14
WT
rnt-1Δlet-7
WT; hsp::bro-1
rnt-1Δlet-7; hsp::bro-1
L2 asymmetric
L3 asymmetric

**Figure S3. Endogenous RNT-1 expression during seam cell development.** (A) Spinning disk images of endogenous RNT-1::GFP expression prior to nuclear envelope breakdown in L1-L3 seam cells. Seam markers are mCherry::PH and mCherry::H2B. Arrowheads point to nuclear RNT-1::GFP. (B) Quantification of the levels of RNT-1::GFP in L1-L3 seam cells. (C) Illustration of the endogenous *rnt-1* 3' untranslated region (3'UTR; red arrowhead) with the *let-7* recognition sites marked in red. These sites are conserved between different nematodes (*C. briggsae, C. remanei, C. brenneri* and *C. japonica*). The red nucleotides were deleted from the endogenous locus using CRISPR/Cas9 recombineering. (D) Spinning disk images of late L4 lateral seam syncytia of control, *rnt-1*$^{\Delta let\text{-}7}$, *hsp::bro-1* and *rnt-1*$^{\Delta let\text{-}7}$ *hsp::bro-1* animals subjected to heat shock prior to the L2 (left panel) and L3 (right panel) asymmetric divisions. (E) Quantification of the number of seam nuclei in late L4 lateral seam syncytia of control, *rnt-1*$^{\Delta let\text{-}7}$, *hsp::bro-1* and *rnt-1*$^{\Delta let\text{-}7}$ *hsp::bro-1* animals subjected to heat shock prior to the L2 and L3 asymmetric divisions. Images were processed using ImageJ software. Scale bars represent 10 µm. Error-bars indicate mean ± SD.

**Figure S4**

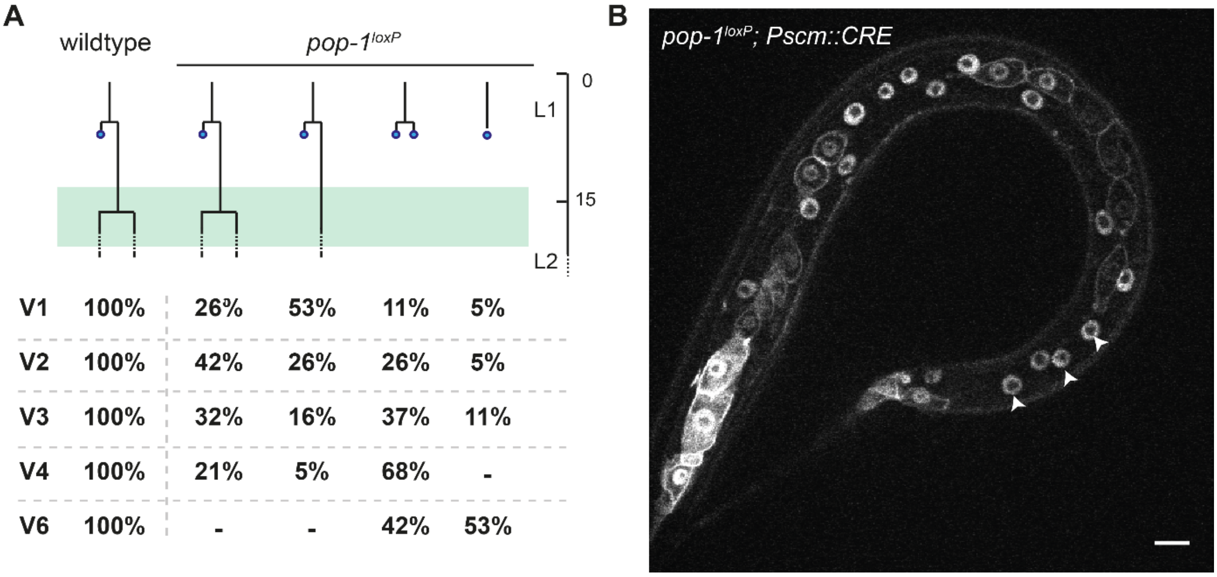


| | wildtype | pop-1loxP | | | |
|---|---|---|---|---|---|
| V1 | 100% | 26% | 53% | 11% | 5% |
| V2 | 100% | 42% | 26% | 26% | 5% |
| V3 | 100% | 32% | 16% | 37% | 11% |
| V4 | 100% | 21% | 5% | 68% | - |
| V6 | 100% | - | - | 42% | 53% |

**Figure S4. Conditional knock-out of *pop-1* reveals a dual role as transcriptional repressor and activator in seam cells.** (A) Lineage analysis of *pop-1$^{loxP}$* early L2 animals undergoing the symmetric seam cell division. Y-axis represents developmental timing (hours; L2 starts at 15 hr). The four different lineages observed in *pop-1$^{loxP}$* animals are plotted against developmental time. The frequency of occurrence of each lineage was quantified for V1-V4 and V6 seam cells. Together, these quantifications reveal extensive premature differentiation of seam cells during L1 stage (B) Spinning disk confocal microscopy images of two late L2 *pop-1*$^{loxP}$, P*scm::CRE* animals. Seam markers are mCherry::PH, mCherry::H2B. Arrowheads points to premature differentiation, asterisk points to extra seam cells. Scale bar represents 10 µm.

**Figure S5**

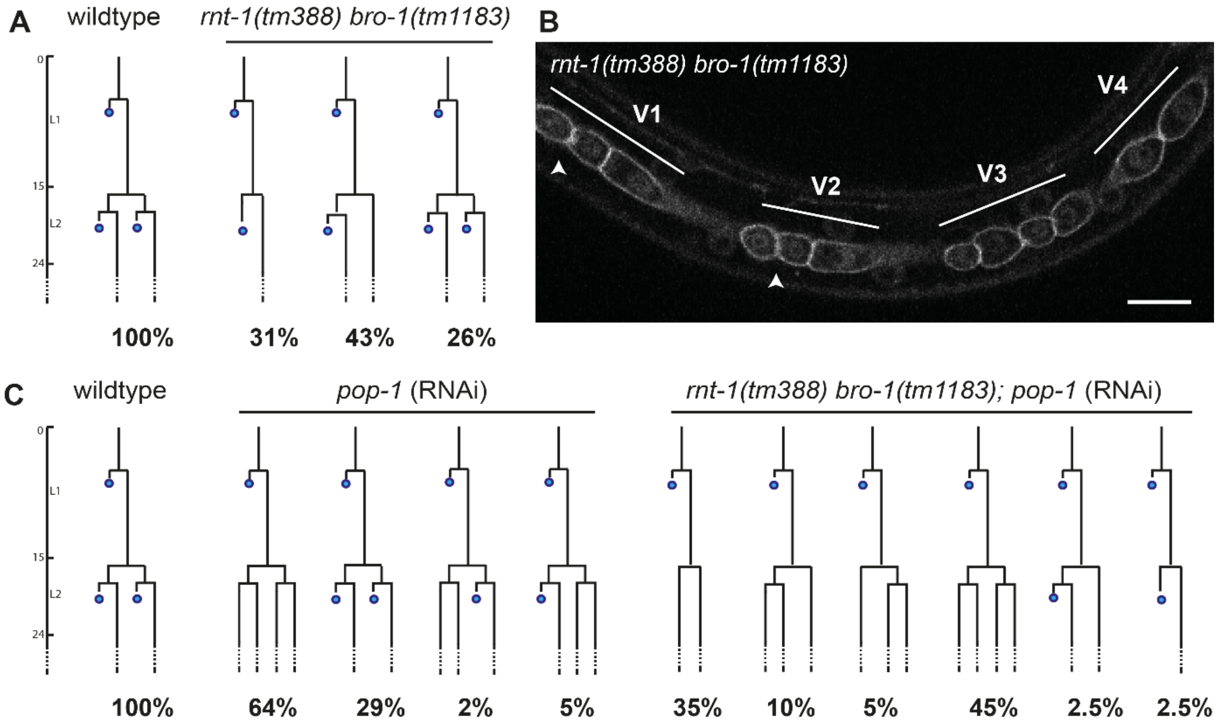


**Figure S5. Overview of lineage analysis of wild-type and mutant strains.** (A) Lineage analysis for wild-type animals and *rnt-1(tm388) bro-1(tm1183)* mutants performed at 20 °C. (B) Spinning disk confocal image of a *rnt-1(tm388) bro-1(tm1183)* animal. The seam markers used are GFP::PH and GFP::H2B. Arrowheads point to anterior daughter cells having undergone the L2 asymmetric division. Scale bar represents 10 µm. (C) Lineage analysis for wild-type animals, *pop-1(RNAi)* animals, and *rnt-1(tm388) bro-1(tm1183)* mutants exposed to *pop-1* RNAi by feeding.

**Figure S6**

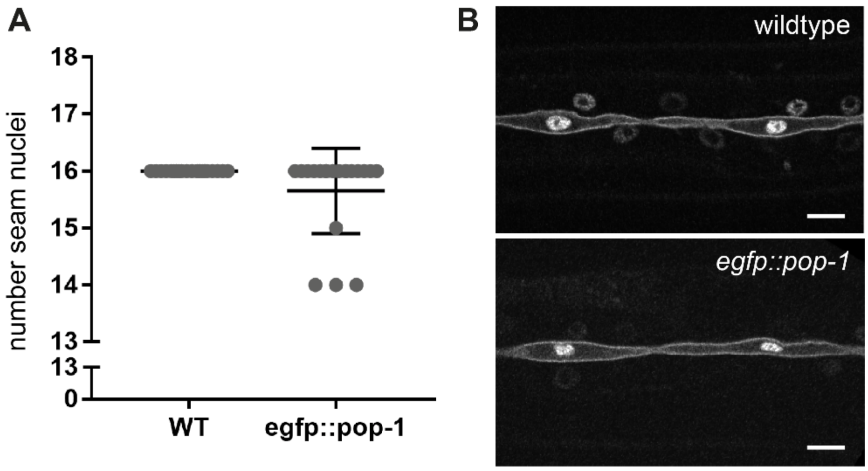


**Figure S6. Characterization of seam epithelia in endogenous *egfp::pop-1* strain.**
Quantification of seam cell nuclei in L4 seam syncytia in wild-type and knock-in animals (A). Representative spinning disk images of L4 stage seam syncytia of wild-type (top) and *egfp::pop-1* (bottom) animals (B). Images were processed with ImageJ software. Scale bars represent 10 µm. Error-bars represent mean ± SD.

**Figure S7**

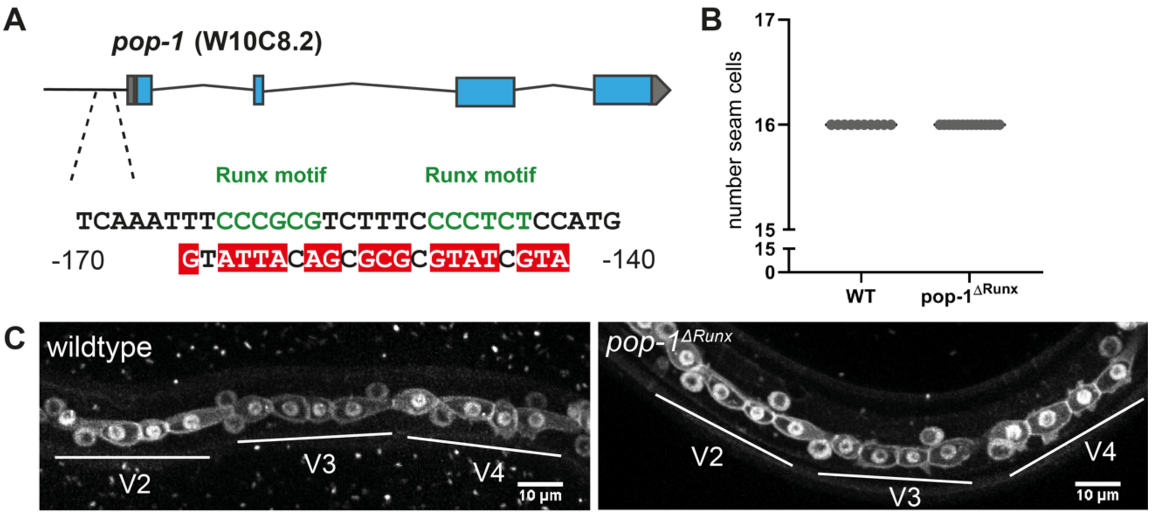


**Figure S7. Mutation of candidate *rnt-1* binding sites in the endogenous *pop-1* promoter region is not sufficient to alter seam cell fate.** Schematic representation of the pop-1 promoter region with the Runx binding motifs. Mutated residues are depicted in red (A). Quantification of seam cell numbers in late L2 animals of *pop1*$^{\Delta Runx}$ animals (B). Representative spinning disk images of *pop1*$^{\Delta Runx}$ late L2 stage seam cells (C). Images were processed using ImageJ software. Scale bars represent 10 µm. Error-bars represent mean ± SD.

### Figure S8

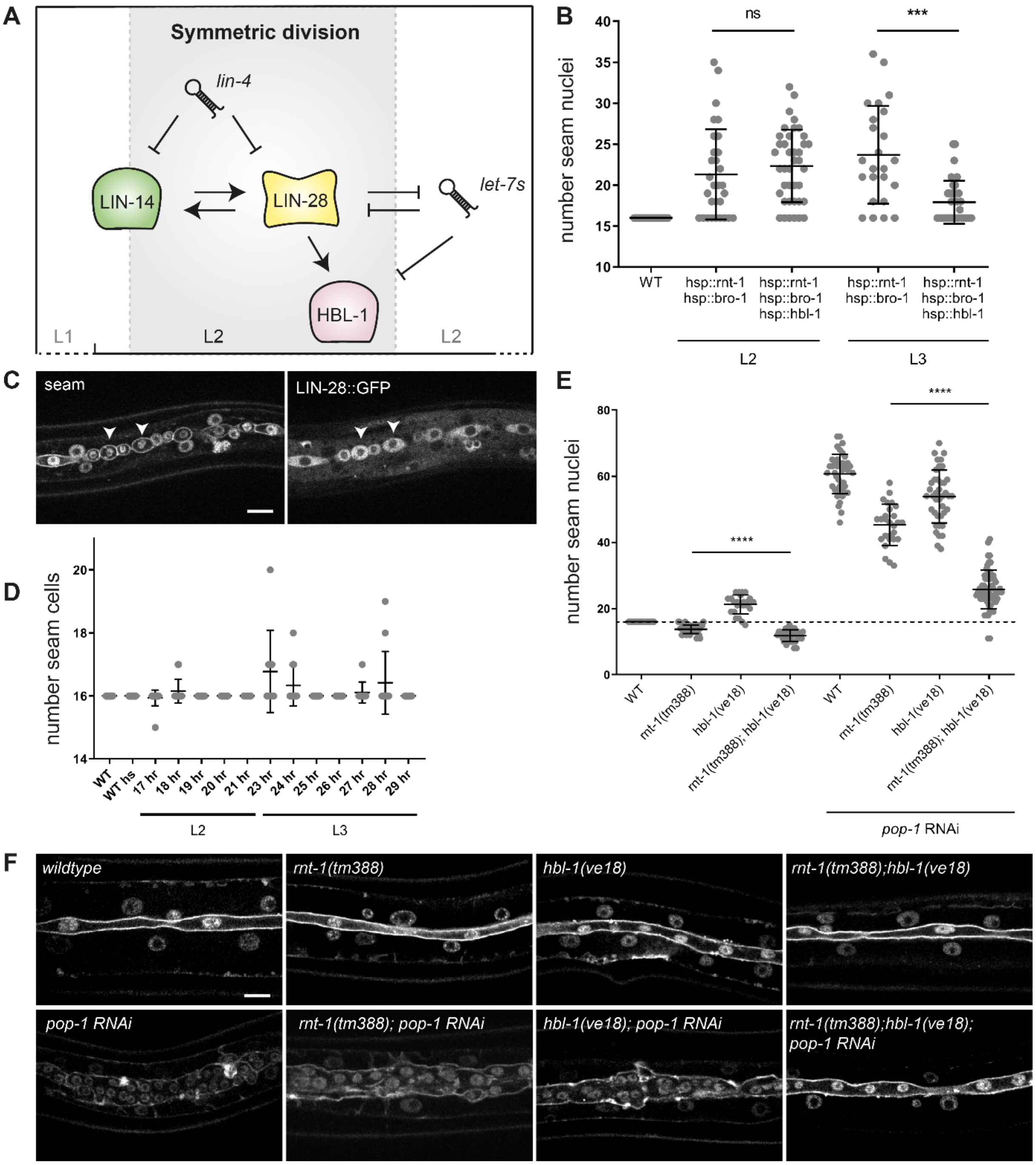


**Figure S8. Manipulation of heterochronic *lin-28* and *hbl-1* gene expression during L2 and L3 asymmetric divisions.** (A) Schematic cartoon of the L2 "symmetry window" as determined by expression of *lin-14*, *lin-28* and *hbl-1* genes. (B) Quantification of the number of seam cell nuclei in late L4 lateral seam syncytia, in animals subjected to heat-shock induction of *rnt-1 bro-1* and *rnt-1 bro-1 hbl-1* prior to the L2 asymmetric or L3 asymmetric division. (C) Spinning disk confocal microscopy image of an L3 larva subjected to heat-shock induction of *lin-28::gfp*. Arrowheads point to posterior daughter cells of the L3 asymmetric seam cell division, displaying high cytoplasmic levels of LIN-28::GFP protein. (D) Quantification of seam cell numbers after heat-shock induction of *lin-28::gfp* in L2 (induction 17 hr-22 hr) or L3 larvae (induction 23-29 hr). Seam cells were counted at the end of L2 (t22 hr) or L3 (t30 hr). (E) Quantification of seam cell nuclei numbers in late L4 lateral seam syncytia of control, *rnt-1(tm388), hbl-1(ve18)* and *rnt-1(tm388);hbl-1(ve18)* animals with and without *pop-1(RNAi)*. (F) Spinning disk confocal microscopy images of late L4 lateral seam syncytia of control, *rnt-1(tm388), hbl-1(ve18)* and *rnt-1(tm388);hbl-1(ve18)* animals with and without *pop-1(RNAi)*. Images were processed using ImageJ software. Scale bars represent 10 µm. Error-bars indicate mean ± SD.

**Table S1. Overview of oligonucleotides used in this study**

| Oligo | Sequence |
|---|---|
| ***pop-1**$^{loxP}$* | |
| *pop-1* N-terminus gRNA 1 | ctcatcgccgagctcttcgtcgg |
| *pop-1* N-terminus gRNA 2 | tttttgtgtatttttatatctgg |
| *pop-1* C-terminus gRNA 1 | taaatgtctactgtagcggaagg |
| *pop-1* C-terminus gRNA 2 | cgctacagtagacatttatgggg |
| *pop-1* N-terminus repair ssDNA oligo | cgttaaaaaatgctctaaatttcaagatataaaaatacacataacttcgtatagcatacattatacgaagttataaaaatgatggcagacgaagagctcggcgatgaggtgaag |
| *pop-1* C-terminus repair ssDNA oligo | ttcatttttctacatcacatgaataacaccataaatgtctataacttcgtatagcatacattatacgaagttatactgtagcggaaagaaaaattaacagcgctacggtagtca |
| *pha-1* repair ssDNA oligo | caaaatacgaatcgaagactcaaaaagagtatgctgtatgattacagatgttcatcaagttattcataaatcattgatag |
| ***egfp::pop-1*** | |
| *pop-1* exon 1 gRNA 1 | gctcatcgccgagctcttcgtcgg |
| *pop-1* LHA FW | ggctgctcttcgtggttggtagaagtctaaacctccacttt |
| *pop-1* LHA RV | gggtgctcttcgcatttttgtgtatttttatatctgg |
| *pop-1* RHA FW | agagctcggcgatgaggtgaaagtgttccgtcgggatgagg |
| *pop-1* RHA RV | gggtgctcttcgtacgaactccgcccataaaaccgt |
| ***pop-1**$^{\Delta Runx}$* | |
| *pop-1* N-terminus gRNA 1 | catggagaggggaaagacgcggg |
| *pop-1* N-terminus gRNA 2 | cggaagttaggccatggagaggg |
| *pop-1* promoter repair oligo | aacctccactttctccccaaaatcctatcgaattcaaatgtattacagcgcgcgtatcgtaatgccctaacttccgcggacctagtccccctttttcttgttttaaatggttcc |
| ***rnt-1::egfp*** | |
| *rnt-1* C-terminus gRNA 1 | atagttcttctccgactatttgg |
| *rnt-1* C-terminus gRNA 2 | gttcttctccgactatttggagg |
| *rnt-1* LHA FW | cagatgccaatgacaatgattccacc |

| | |
|---|---|
| *rnt-1* LHA RV | aaaaggtctccatatcgtaggtgatgagctattcgatgaagt |
| *rnt-1* RHA FW | tcttaaaaatattcattattttaccacaacacacc |
| *rnt-1* RHA RV | tctaatcatccatctcccaactc |
| **Heat shock expression** | |
| FW *hsp16.48* | ctggacggaaatagtggtaaag |
| RV *hsp16.48* | tcttgaagtttagagaatgaacag |
| FW *unc-54 UTR* | catctcgcgcccgtgcc |
| RV *unc-54 UTR* | aaacagttatgtttggtatattggg |
| FW *cki-1* | atgtcttctgctcgtcgttg |
| RV *cki-1* | gtatggagagcatgaagatcg |
| FW *egfp* | atgtccaagggagaggagc |
| RV *egfp* | ttacttgtagagctcgtccattcc |
